# An Allee-based Distributed Algorithm for Microbial Whole-Cell Sensors

**DOI:** 10.1101/2023.08.25.554781

**Authors:** Fabricio Cravo, Matthias Függer, Thomas Nowak

## Abstract

Reliable detection of substances present at potentially low concentrations is a problem common to many biomedical applications. Complementary to well-established enzyme-, antibody-antigen-, and sequencing-based approaches, so-called microbial whole-cell sensors, i.e., synthetically engineered microbial cells that sense and report substances, have been proposed as alternatives. Typically these cells operate independently: a cell reports an analyte upon local detection.

In this work, we analyze a distributed algorithm for microbial whole-cell sensors, where cells communicate to coordinate if an analyte has been detected. The algorithm, inspired by the Allee effect in biological populations, causes cells to alternate between a logical 0 and 1 state in response to reacting with the particle of interest. When the cells in the logical 1 state exceed a threshold, the algorithm converts the remaining cells to the logical 1 state, representing an easily-detectable output signal. We validate the algorithm through mathematical analysis and simulations, demonstrating that it works correctly even in noisy cellular environments.

## 1 Introduction

Numerous disease indicators are based on detecting that the abundance of a particular substance exceeds a threshold concentration [3, 23, 26]. Widely-adapted techniques are sequencing for genetic information and antibody-based detection for proteins [20]. Recently microbial whole-cell sensors (MWCS), i.e., cells engineered to sense and report substances, emerge as an easy-to-use and cost-effective alternative to these classical detection methods [2]. MWCS have been demonstrated to successfully sense pollutants [4], detect inflammation in mice models [24], and provide means for environmental monitoring [16, 12, 19].

### Local and population-based sensors

To detect analytes at low concentrations, MWCS use techniques such as optimizing the cellular sensing circuitry [13, 24] and sensing multiple, correlated analytes [24]. These approaches are examples for *local sensor designs*, where engineered cells locally sense and report analytes. In this case, the population-level readout is obtained as the cumulative single-cell responses, which inherently limits the population-level response: Assume a population *C* of *n* cells, a small fraction *α* ∈ [0, 1] of which detect the analyte of interest, and an ideal local cell response out_*c*_(in_*c*_) that maps the presence (in_*c*_ = 1) or absence (in_*c*_ = 0) of a detection event at cell *c* to a local cell output. In presence of an ideal local cell response that outputs a (normalized) 1 if in_*c*_ = 1 and 0 otherwise, the population-level response out_pop_ is given by

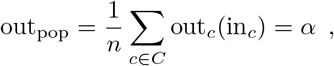

i.e., remains linear in the fraction *α* of cells that detect the analyte.

By contrast, *population-based designs* use communication between cells to potentially achieve improved threshold-like population responses. In terms of the example before this is achieved by allowing out_*c*_ to depend not only on in_*c*_, but also on the other cells’ (communicated) inputs: the population-level response

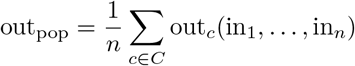

is not necessarily proportional to *α* in this case.

A commonly used mechanism to communicate is via quorum sensing (QS) molecules [25, 18], which diffuse through the cell membrane and, if present in sufficiently high concentrations, allow cells to trigger a population-level response [16, 12, 19]. For example, the circuit by Hsu, Chen, Hu, and Chen [14] uses QS for population-level signal amplification: when cells sense metal ions, they start to secrete the QS molecule. This molecule, then diffuses into the surrounding medium and inside the population’s cells. If a cell’s internal concentration exceeds a certain threshold, a reporting pathway is triggered. Further examples for population-based designs based on QS are the detection of mercury [6] and phenolic compounds [13].

### Allee-based algorithm

In this article, we analyze a simple distributed algorithm that acts as a distributed amplification circuit and, together with a local sensory and reporting circuit, yields a population-based MWCS design. In our algorithm, cells transition from a low (L) state to a high (H) state upon reacting with rare-event substances of interest. When a specific number of cells successfully alter their state within a predetermined time frame, the algorithm guarantees the production of an amplified reporter signal, indicating the presence of rare events. Conversely, if this threshold is not met, the reporter signal is guaranteed to remain low and is not amplified at population-level. Given the high noise levels in biological circuits [15], the threshold is designed to mitigate the number of false positives compared to naive broadcasting methods, offering noise protection.

The algorithm is inspired by the Allee effect [1] as observed in biological populations: While in populations that compete for a shared resource, lower densities are supposedly more likely to thrive [8], the Allee effect describes the phenomenon that the fitness of small populations often decreases, e.g., due to the reliance on cooperation strategies within the population [5, 17, 7]. The algorithm is designed such that the population of H cells shows an Allee-like behavior: for low H densities, the “birth” rate, i.e., the rate by which L cells are transformed into H cells, is compensated by their “death” rate, i.e., the rate by which H cells are transformed into L cells. Above a certain threshold cell density, the situation is reversed, and the birth rate outweighs the death rate.

Further, differently than other QS amplification circuits [13, 14, 6], our algorithm autonomously maintains the amplified state indefinitely. This is achieved through a positive feedback loop, where the signal responsible for amplification promotes itself.

### Organization

We define the model and algorithm as a reaction network in Section 2. Section 3 establishes correctness conditions (Theorem 1) and a bound for the time necessary to trigger the amplified output signal (Theorem 2). We provide simulations for a genetic circuit implementation in Section 4, compare its properties to a naive solution in Section 5, and assess the algorithm’s robustness to noise in Section 6. A discussion in Section 7 concludes the work.

## 2 Model and Algorithm

The concentration of the analyte of interest is denoted by *A*(*t*) ∈ ℝ_+_. Cells of the detection algorithm are in either of two states: low (L), voting for the absence of the analyte of interest, or high (H), voting for its presence. We denote the density of cells in L at time *t* by *L*(*t*) ∈ ℝ_+_, and the density of cells in H by *H*(*t*) ∈ ℝ_+_. We write *P* (*t*) = *H*(*t*) + *L*(*t*) for the total population size. Assuming that the population size is within a steady state, we neglect replication and cell-death reactions.

### A broadcasting algorithm

A naive algorithm to obtain a non-linear population-level response would be to broadcast any detection of the analyte by a cell to all other cells. Such an algorithm comprises of two reactions

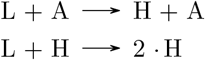

the first of which models detection of an analyte by a cell and the second the broadcast of such an event to all other cells. However, as we show in Section 5, this algorithm does not tolerate incorrect detections, i.e., cells that incorrectly transition from state L to state H in absence of an analyte; a problem any biological implementation will necessarily have.

### The Allee-based algorithm

To address the problem of the broadcasting algorithm to deal with faulty state transitions, we propose an algorithm that tolerates erroneous detection of the analyte up to a certain rate. Our algorithm comprises of three reactions that determine when a cell switches state: (i) Reaction (Detect): A cell in state L changes to state H upon local detection of the analyte. We assume that this happens with a rate *σ*_*A*_ ∈ ℝ_+_. (ii) Reaction (Hold): A cell in state L also switches to state H with a rate that depends on the density *H* according to a Hill function with parameters *κ* ∈ ℝ_*>*0_, *K* ∈ ℝ_*>*0_, and *n* ∈ ℝ with *n >* 1. The Hill function models the fact that this reaction is triggered by a QS molecule which is secreted by cells in the H state; see, e.g., [11] for a Hill-function model of a QS circuit. (iii) Reaction (Reset): A cell in state H switches back to state L with a certain *reset rate ρ* ∈ ℝ_*>*0_. Intuitively, this is to prevent accumulation of incorrect detection events in the system.

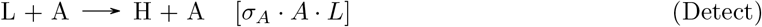

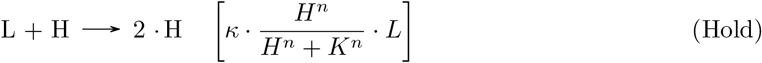

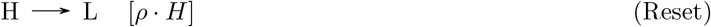

Cells may incorrectly detect the analyte with a rate *σ*_err_ ∈ ℝ_*>*0_, accounted for in the additionally reaction

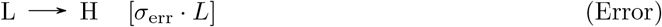

For the purpose of analysis, unless stated otherwise, we will subsume both (Detect) and (Error) into the single reaction

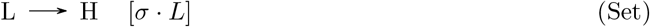

calling *σ* = *σ*_*A*_ · *A* + *σ*_err_ the *rare-event detection rate*.

Figure 1 illustrates the algorithm’s reactions (Figure 1a) and three modes of operation: in absence of the analyte (Figure 1b), in its presence (Figure 1c), and after detection of the analyte with the analyte potentially being absent (Figure 1d). Observe that the high density of H cells is maintained in the latter case.

**Figure 1:**
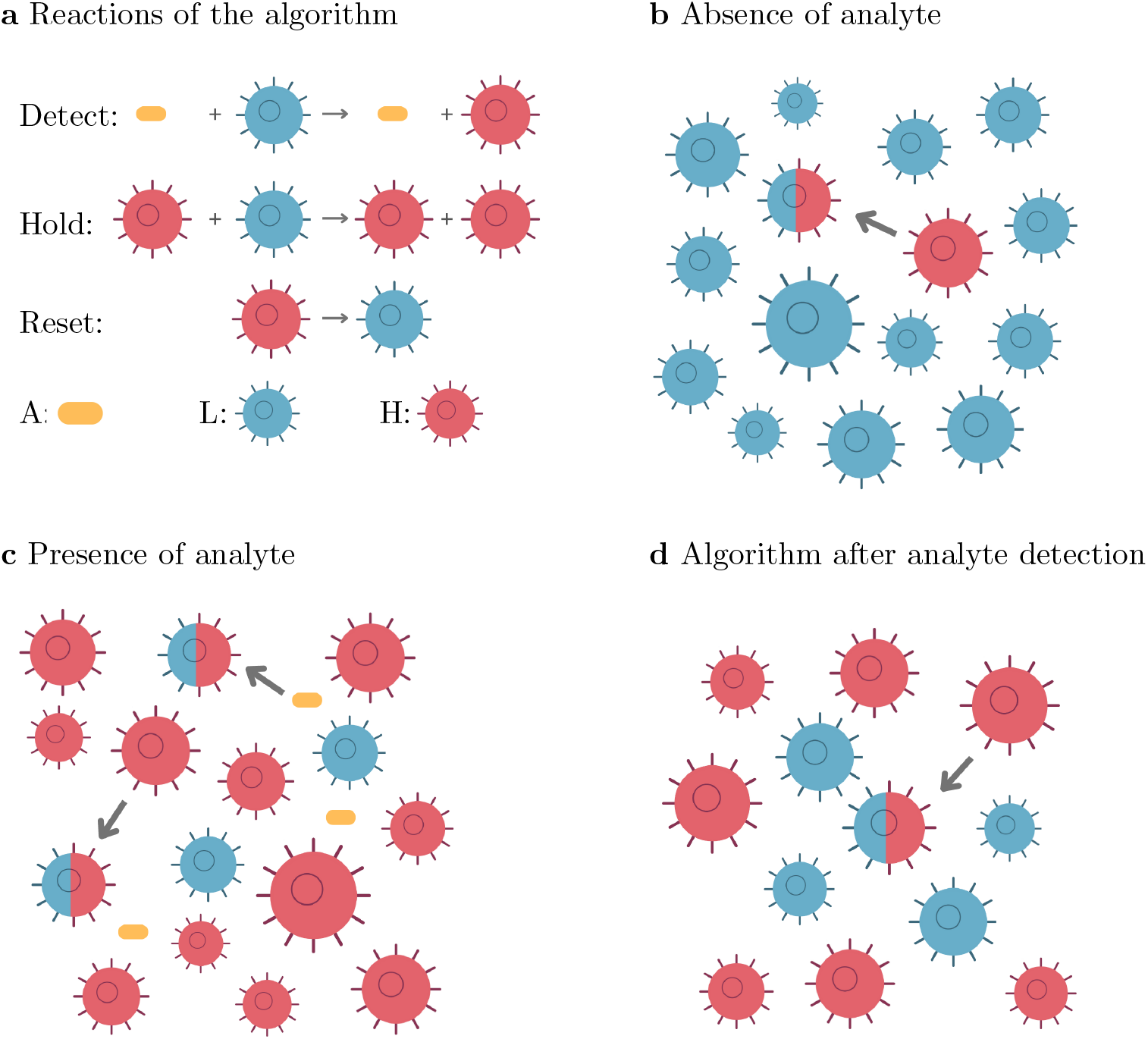
Illustration of the algorithm: **a** The three reactions (Detect), (Hold), and (Reset). **b** Cell population in absence of the analyte. A small amount of cells are incorrectly converted from L to H, but the population remains mostly in the L state. **c** Cell population in presence of a sufficiently high concentration of the analyte. Interaction with the analyte converts L cells into H cells. Additionally H cells convert other L cells into H cells. **d** State of the population after the analyte has been detected and was potentially removed thereafter. The high density of H cells is maintained by H cells continuously converting L cells into H cells.

Figure 2 visualizes the Allee-like behavior of H cells: Figure 2a&b show the “birth” rate of H cells, i.e., the sum of the rates in (Hold) and in (Set), as well as the “death” rate of H cells, i.e., the rate in (Reset), over the density of H cells. In the absence of the analyte (Figure 2a), the rare-event detection rate *σ* is low, and stable steady states for the H density are either low (close to 0 mL^−1^) or high (about 1.5 · 10^8^ mL^−1^). We show in Section 3 that the low-density steady state is reached if the initial H density was below a threshold, and the high-density steady state if it was above this threshold, thus guaranteeing the memory effect for a once detected analyte. Conversely, in presence of the analyte, and a consequently high rare event detection rate *σ*, the H cell density converges to a high value (Figure 2b).

**Figure 2:**
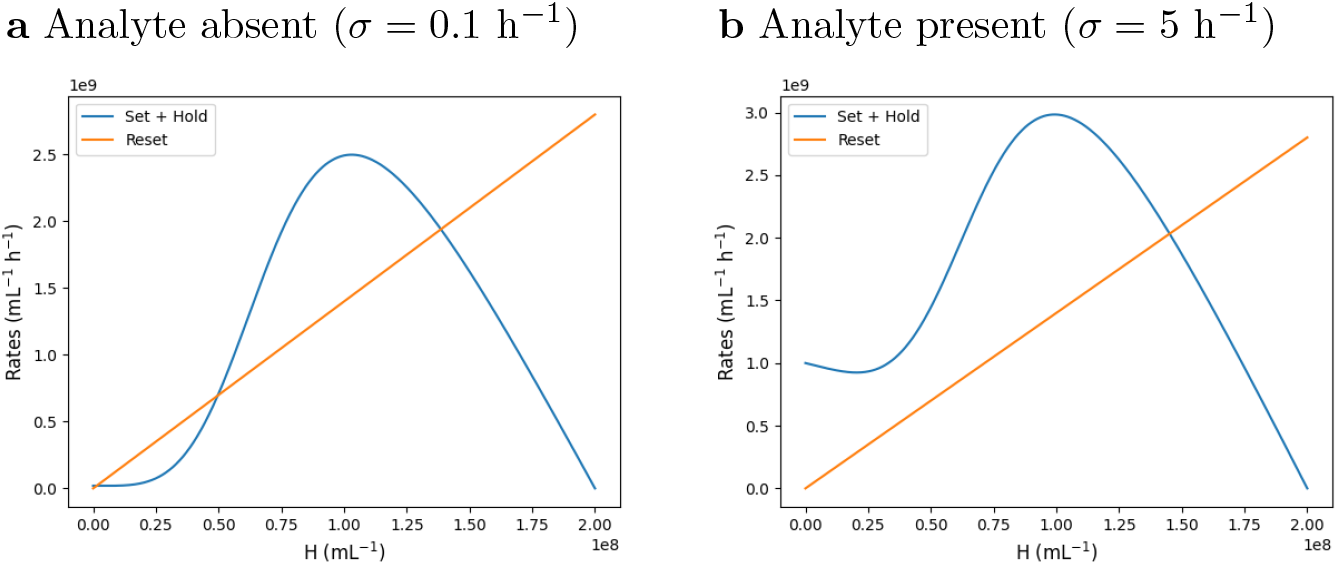
Allee-like behavior of the H cells. **a&b** Birth rate (Set) + (Hold) of H cells (orange) and death rate (Detect) of H cells (blue) versus H density. The characteristic s-shape of a birth rate in a population with Allee effect is visible. **a** Rates in absence of the analyte (*σ* = 0.1 h^−1^). **b** Rates in presence of the analyte (*σ* = 5 h^−1^). Parameters used: *κ* = 35 h^−1^, *ρ* = 14 h^−1^, *P* = 2 10^8^ mL^−1^, *n* = 4

## 3 Correctness of the Algorithm

We can write an ordinary differential equation (ODE) for the cell densities *H* and *P* from reactions (Set), (Reset), and (Hold), obtaining

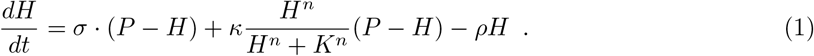

Algebraic manipulation of 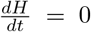 yields up to three potential equilibrium points for *H*, the smallest of which is stable, the middle one unstable, and the largest one stable (if they exist). Using the monotonicity of 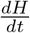 within certain subdomains of *H*, one can show that for the interval *I* between the first, stable, and the second, unstable, equilibrium point, 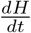 is negative for *H* ∈ *I*, and *I* = ∅ for rare event detection rates *σ* larger than a critical rate *σ*_*c*_. The proof we give in the Supplementary Material (Section A) is based on the intermediate value theorem. For sufficiently large values of *σ*, only one stable equilibrium point remains and its value can be bounded away from 0. The proof in the Supplementary Material uses a quadratic Lyapunov function to show convergence to this fixed point.

Based on this we can establish the correctness of the Allee-based algorithm by showing that under certain conditions of its parameters *κ, K, n*, and the total population size *P*, the algorithm guarantees: (i) convergence to a low density of H cells if the rare event detection rate *σ* is below a critical rate *σ*_*c*_ and the initial density of H cells is low (no memorized detection happened), and (ii) to a high density of H cells either if the initial population of H cells was high (a detection was memorized), or the rare event rate *σ* exceeds the critical threshold rate *σ*_*c*_ (the analyte is being detected).

### Theorem 1.

*If* 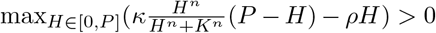, *then there exist α*_*i*_, *α*_*f*_ *in*]0, 1[*with α*_*i*_ *< α*_*f*_ *and σ*_*c*_ *>* 0 *such that:*

- *If σ < σ*_*c*_, *there exists a critical point H*_*c,σ*_ *such that:*
  1. *If H*(0) *< H*_*c,σ*_, *then H*(*t*) *converges to a value in* [0, *α*_*i*_*P* [.
  2. *If H*(0) *> H*_*c,σ*_, *then H*(*t*) *converges to a value in* [*α*_*f*_ *P, P* [.
- *If σ > σ*_*c*_, *then H*(*t*) *converges to a value in*]*α*_*f*_ *P, P* [.

The following two corollaries immediately follow from Theorem 1 and establish the correctness of detection (Corollary 1) and memorization of a previously detected analyte (Corollary 2).

### Corollary 1 (Detection)

*If the conditions for Theorem 1 hold, and with α*_*i*_, *α*_*f*_, *and σ*_*c*_ *as defined in Theorem 1, if H*(0) = 0 *then H*(*t*) *converges to a value in* [0, *α*_*i*_*P* [*if σ < σ*_*c*_, *and to a value in* [*α*_*f*_ *P, P* [*if σ > σ*_*c*_.

### Corollary 2 (Memory)

*If the conditions for Theorem 1 hold, and under the notation for Theorem 1, if H*(0) *> α*_*i*_*P, then H*(*t*) *converges to a value in* [*α*_*f*_ *P, P* [.

We finally establish an upper bound on the time the algorithm needs to converge to a high density of H cells in presence of an analyte. The proof is given in the Supplementary Material (Section A) and analyzes a bounding ODE 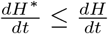, for which the convergence time is established.

### Theorem 2 (Convergence time)

*Let σ > σ*_*c*_. *Let* 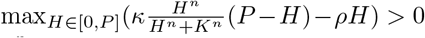. *Let H*_*c*,0_ *be the second lowest non-negative root solution of* 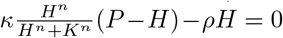. *If* 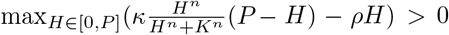, *there exists a time* 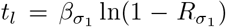 *with β*_*σ*_ *and R*_*σ*_ *functions of σ, such that H*(*t*) *> H*_*c*,0_ *for any t > t*_*l*_.

## 4 Simulation Results

To validate that the algorithm performs well within realistic parameter ranges, we estimated parameters and ran simulations from a potential genetic circuit implementation (Figure 4). Following previous QS circuit designs in synthetic biology [10, 11], we use an N-acyl homoserine lactone (AHL) as the QS molecule. The AHL molecule is synthesized by LuxI under the control of a promoter (p1 in Figure 4) that is activated by the binding of an LuxR-AHL complex. LuxR is consituently expressed by the circuit (not shown in the figure). Additionally, the detection of the analyte by the cell is assumed to activate promoter p1.

**Figure 3:**
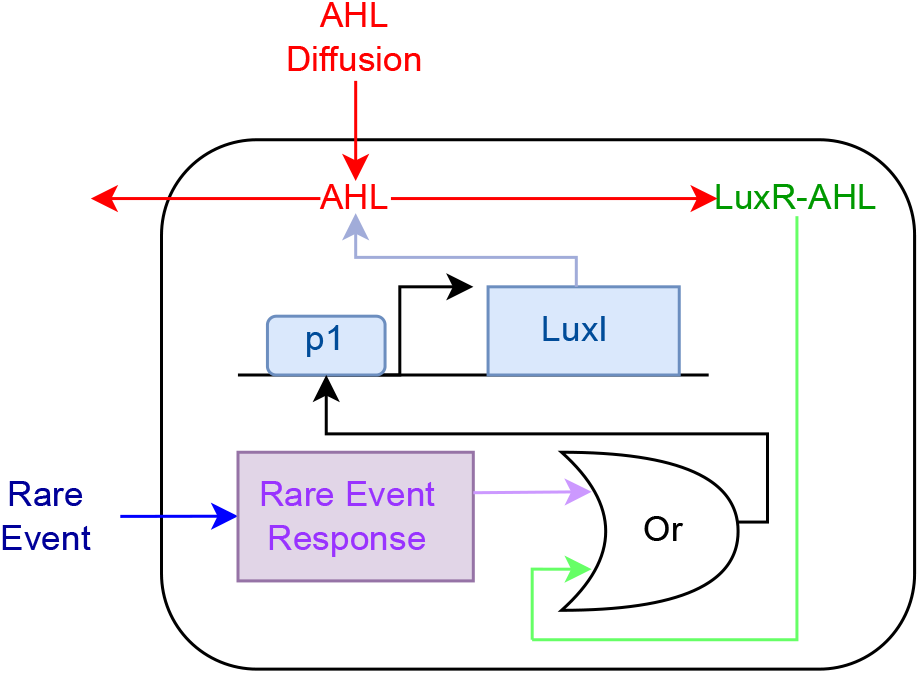
Genetic circuit implementation of the Allee-based algorithm. Both the AHL and the rare event can transform the cell from the state L to H by triggering the expression of LuxI. LuxR is constitutively expressed (circuit not shown).

**Figure 4:**
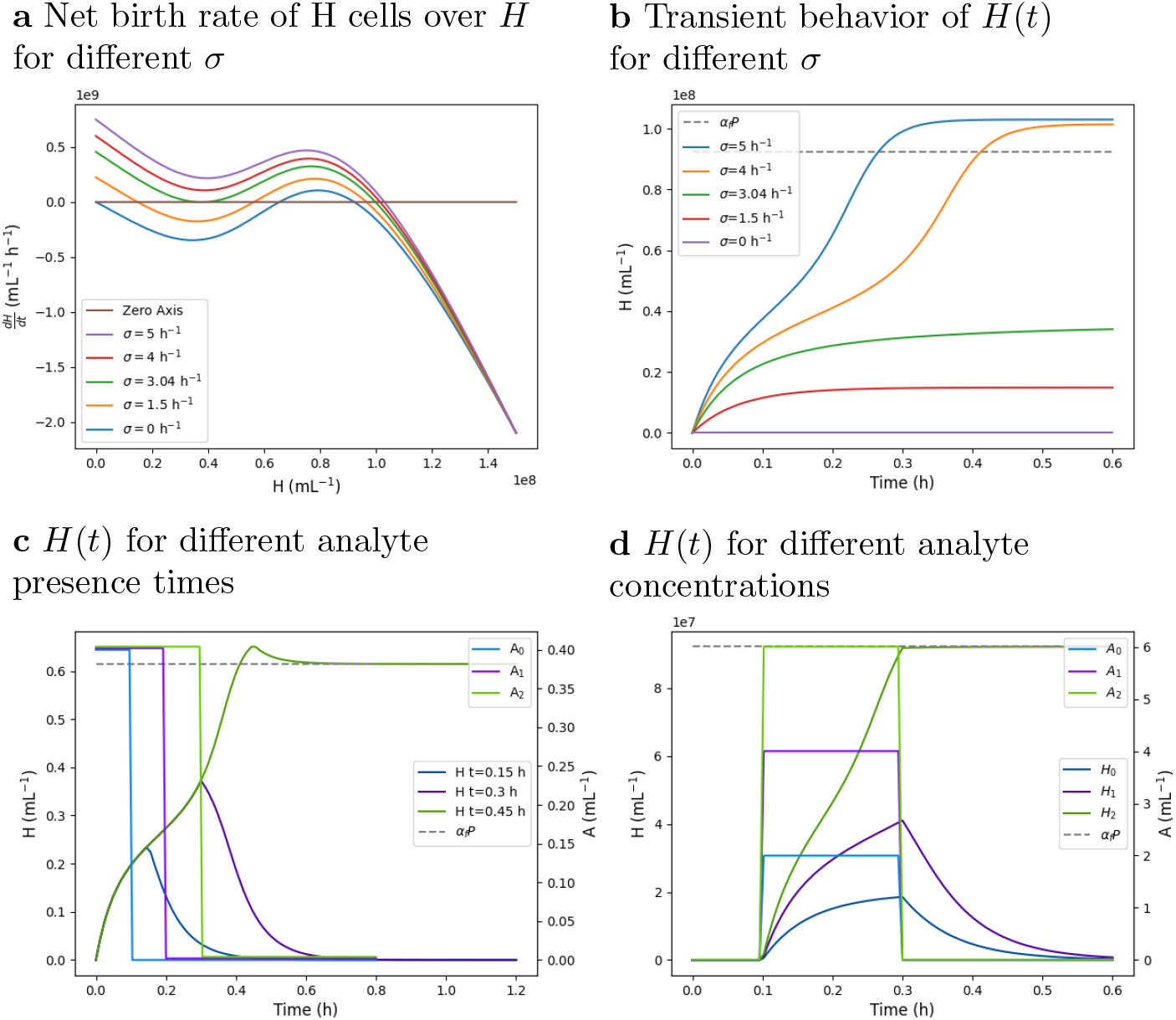
Simulation results. **a** Plot of the net birth rate H cells (*dH/dt*) versus *H* for different *σ*. Equilibrium points of *H* are the roots of *dH/dt*. **b** Plot of *H*(*t*) over time *t*. For values of *σ* bellow the critical *σ*_*c*_, *H*(*t*) converges to H cell densities below *α*_*i*_*P* (consistent with Corollary 1). For values of *σ* above *σ*_*c*_, *H*(*t*) converges to H cell densities above *α*_*f*_ *P* (again, consistent with Corollary 1). For *σ* = 3.05 *h*^−1^, the plotted time range is not enough to show convergence of *H*(*t*). **c** Plot of *H*(*t*) and the analyte concentration *A*(*t*) over time *t*. The analyte concentration was chosen as a negative step function with different durations until the step. One observes that below a certain exposure time of the cells to the analyte, the analyte is not memorized by the algorithm. Above this exposure time, the detection of the analyte is memorized even if it removed thereafter (consistent with Corollary 2). **d** Plot of *H*(*t*) and the analyte concentration *A*(*t*) over time *t*. The analyte concentration was chosen as a pulse of varying amplitude. One observes that below a certain amplitude, the detection is not memorized, while above it is memorized.

We next outline how this circuit implements the Allee-based algorithm: The algorithm’s cell states L and H model cells with low internal LuxI, respectively, high internal LuxI concentrations. Reaction (Set) models the fact that an anlyte leads to an increasing internal concentration of LuxI, thus converting an L cell to an H cell. Also, LuxI is degraded and diluted within the cell, accounting for reaction (Reset). Finally, H cells synthesize AHL that diffuses into the medium and from there into surrounding cells. The so-formed LuxR-AHL complex consequently activates promoter p1 that shows a Hill-type activation profile. The promoter’s activation again leads to expression of LuxI, making an L cell switch to an H cell, as required by reaction (Hold).

### Reaction and population parameters

For simulations we parametrized the algorithm’s reactions with rate parameters from literature (Table 1). For the previously discussed implementation, *κ* corresponds to the expression of LuxI controlled by p1. From the model by Din *et al*. [11, supplementary information], we choose *κ* = 35 h^−1^. The reaction rate constant *ρ* corresponds to the degradation rate constant of LuxI and was set to 14 h^−1^ [11]. The Hill coefficient *n* for activation of p1 via LuxR-AHL was set to 4 [11]. The threshold parameter *K* for the activation via LuxR-AHL was set to a relatively high value of 8 · 10^7^ mL^−1^, reported in a circuit by Smith and Schuster [21]. The total cell density was set to a value larger than 2-times the threshold *K*, and well in the range of reachable *E. coli* cell densities; we used *P* = 1.5 · 10^8^ mL^−1^. The parameters in Table 1 fulfill the condition of Theorem 1.

**Table 1:** Reaction and population parameters used in simulations.

| Parameter | Value | Source |
| --- | --- | --- |
| $\kappa$ | $35 \text{ h}^{-1}$ | [11] |
| $\rho$ | $14 \text{ h}^{-1}$ | [11] |
| $n$ | 4 | [11] |
| $K$ | $8 \cdot 10^7 \text{ mL}^{-1}$ | [21] |
| $P$ | $1.5 \cdot 10^8 \text{ mL}^{-1}$ | assumed, $> 2K$ |

The critical threshold rate *σ*_*c*_ was determined through binary search. We start from the interval *I* = [0, *σ*_utr_], where 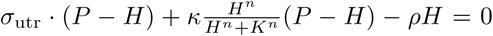 has only one solution for *H*. From Theorem 1, the equation 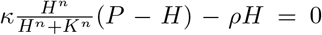 necessarily has three solutions for *H*. We then repeatedly determine the midpoint of the interval, and verify if the above equation has three or one solution with the midpoint as *σ*. In case of three solutions, we replace the left bound of *I* by the midpoint, and in case of one solution we replace the right bound by the midpoint. In case two solutions or a sufficient precision is reached, the search terminates. We refer the reader to the Supplementary Material for more details. For our setting, we find that *σ*_*c*_ *≈* 3.04 h^−1^, about 22% of the system’s lowest rate constant, which is *ρ*.

### Simulation of the algorithm

We used the Python simulation framework MobsPy [9] to obtain transient and steady-state simulation results for the parameters in Table 1. The core of the simulation code is shown in Listing 1.

**Listing 1:**
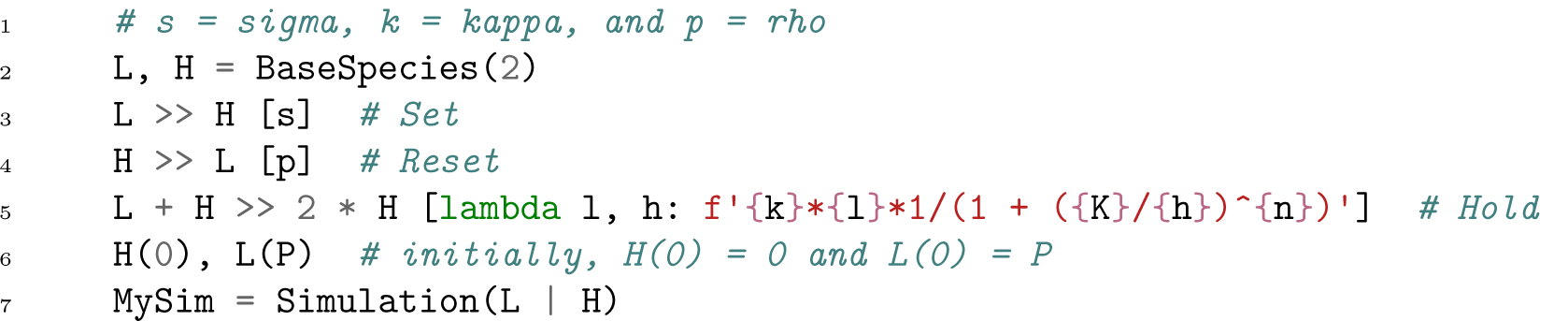
MobsPy simulation code for the Alle-effect-based algorithm. Initialization of reaction and population parameters is according to Table 1. Parameter *σ* (s in the code) was varied in the simulations.

Figure 4a shows 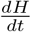, i.e., the net birth rate of H cells, versus *H* for different rare event detection rates *σ*. While larger values of *σ* lead to a single equilibrium points, lower values (0 h^−1^ and 1.5 h^−1^ in the figure) result in three equilibrium points. Of the three points, the smallest and the largest are stable and are seen to have different *H* densities for different *σ*: the smallest equilibrium point corresponds to a negative detection result and the largest to a positive detection result. Further, choosing *σ* = 3.04 h^−1^ close to the critical *σ*_*c*_, results in a net birth rate function that barely touches the x-axis.

We next ran transient simulations for the same *σ* values as in Figure 4a over a time range of 0.6 h simulated time (Figure 4b). One observes, the convergence of *H*(*t*) to a high density of H cells for a *σ* of 4 h^−1^ and 5 h^−1^, and to a low density of H cells for a *σ* of 0 h^−1^ and 1.5 h^−1^; in agreement with Theorem 1. The transient simulation for *σ* = 3.04 h^−1^ close to the critical *σ*_*c*_ does not visibly converge in the simulated time. Increasing the simulation time, however, shows that it converges to a high H density.

Figure 4c&d demonstrate the memorization of the presence of an analyte that has been removed thereafter. In Figure 4c we varied the exposure time of the cells to the analyte, while in Figure 4d the concentration of the analyte was varied. The memorization above a certain critical rate *σ*_*c*_(*t*) = *σ*_*A*_ · *A*(*t*) is observed in both cases, which is consistent with Corollary 2.

We finally ran simulations for an extended duration to determine steady-state values for different settings of rare event detection rates *σ* (Table 2). The results are consistent with Theorem 1. When *σ* is above the critical rate *σ*_*c*_, larger values of *σ* lead to faster convergence of *H*(*t*) to its steady state. As shown in Table 2, the highest convergence time is 0.61 h, which occurs for *σ* = 3.04 h^−1^, a value close to *σ*_*c*_. For a slightly higher *σ* = 4 h^−1^, the convergence time is already reduced to 0.44 h. For a mathematical analysis of the convergence times, we refer the reader to the Supplementary Material (Lemmas 7 and 19).

**Table 2:** Steady-state density of *H*(*t*) and convergence times for different rare event detection rates *σ*. Convergence times are given as the time until 95% of the steady-state density is reached.

| Rate $\sigma$ | Steady-state of $H$ | Convergence time |
| --- | --- | --- |
| $0.00 \text{ h}^{-1}$ | $0 \text{ mL}^{-1}$ | 0 h |
| $1.50 \text{ h}^{-1}$ | $0.15 \cdot 10^8 \text{ mL}^{-1}$ | 0.21 h |
| $3.04 \text{ h}^{-1}$ | $0.36 \cdot 10^8 \text{ mL}^{-1}$ | 0.61 h |
| $4.00 \text{ h}^{-1}$ | $1.01 \cdot 10^8 \text{ mL}^{-1}$ | 0.44 h |
| $5.00 \text{ h}^{-1}$ | $1.03 \cdot 10^8 \text{ mL}^{-1}$ | 0.3 h |

## 5 Comparison with Other Algorithms

To demonstrate the effectiveness of the Allee-based algorithm, we compare its performance to other algorithms, including two natural adaptations and one presented by Hsu, Chen, Hu, and Chen [14]. In the comparison we focus on the thresholding behavior of the algorithms: ideally the detection algorithm shows a strong amplification of its detection output around a threshold concentration of the analyte, below of which the output is strong negative, and above of which it is strong positive.

### Set-reset algorithm

A natural simplification of the Allee-based algorithm is to remove the (Hold) reaction, and only keep the reactions that transform cells to H cells in presence of the analyte, as well as reset H cells to L cells with a certain reset rate *ρ*:

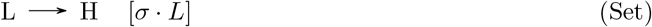

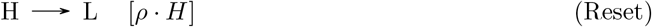

Steady-state analysis of H cells via setting 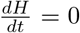 and subsequent algebraic manipulation yields, 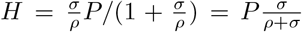 as the unique steady-state. Since it is unique, the algorithm lacks the possibility to memorize previous presence of the analyte. Further, for most applications we expect *σ* to be small compared to the other rates (and in particular *ρ*), implying a low amplification from the analyte concentration *A* to the output *H*.

### Broadcasting algorithm

A natural distributed algorithm that solves the problem of detecting an analyte is to broadcast any detection of the analyte to all other cells that relay this broadcast. Here, relay is obtained by a cell in state H that had been informed of the presence of the analyte, to pass this information to any L cell it interacts with. In terms of reactions, and referring to *κ >* 0 as the broadcasting rate, this algorithm can be written as

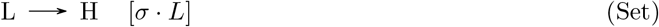

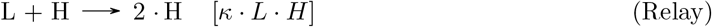

and its dynamics are 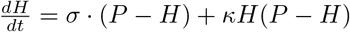.

Since 0 ≤ *H* ≤ *P*, one has that *H* is bounded. We now perform a case distinction between the cases where *σ >* 0 and *σ* = 0. **Case** *σ* = 0: For any *H* ∈ [0, *P* [, one has 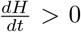. From the monotonicity of *H* and its boundedness, it follows that *H* converges to a finite steady-state. Since 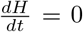 when *H* = *P*, one has that *H*(*t*) converges to *P*. **Case** *σ >* 0: For any *H*]0, *P* [, one has 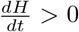. If *H*(0) is in]0, *P*], from the monotonicity of *H* in]0, *P*] and its boundedness, it follows that *H* converges to a finite steady-state. Since 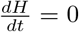 when *H* = *P*, one has that *H*(*t*) converges to *P*. If *H*(0) = 0, since 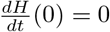, one has that *H*(*t*) = 0 for all *t*. Indeed, two steady states, obtained by setting 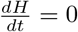, are possible: (i) *σ* = 0 and *H* = 0, and (ii) *σ >* 0 and *H* = *P*.

While the presence of the two steady-states shows that the algorithm can memorize previously detected analytes, the steady-states also show that the algorithm cannot tolerate incorrect transitions of L cells to H cells: for an arbitrarily small *σ*, all cells switch to state H.

### Distributed amplification algorithm

Hsu, Chen, Hu, and Chen [14] proposed a circuit that uses distributed amplification via a QS pathway: Cells that detect the analyte synthesize the QS molecule. Similar to the model for the Allee-based algorithm, we abstract this via two cell types: L cells with low internal concentrations of LuxI and H cells with high concentrations of LuxI. Any cell whose QS threshold is triggered, expresses a reporter molecule S (e.g., YTP). In terms of reactions, this algorithm is expressed as

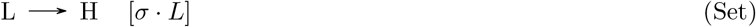

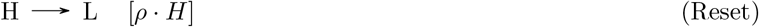

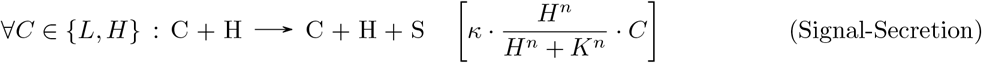

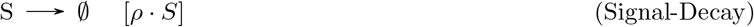

where S is the reporter molecule, and the reaction (Signal-Decay) accounts for the decay of S.

To compare the thresholding behavior of the distributed amplification, the Set-reset, and the broadcasting algorithm to the Allee-based algorithm, we ran simulations in presence of the same analyte concentration *A* for all four algorithms (Figure 5). Simulation parameters where chosen identically for similar reactions. One observes the strong amplification of the Allee-based algorithm around a non-zero critical concentration of A. The distributed amplification algorithm shows a threshold behavior, but with a weaker amplification. The broadcasting algorithm has a strong threshold at 0, and the Set-reset algorithm shows no thresholding behavior.

**Figure 5:**
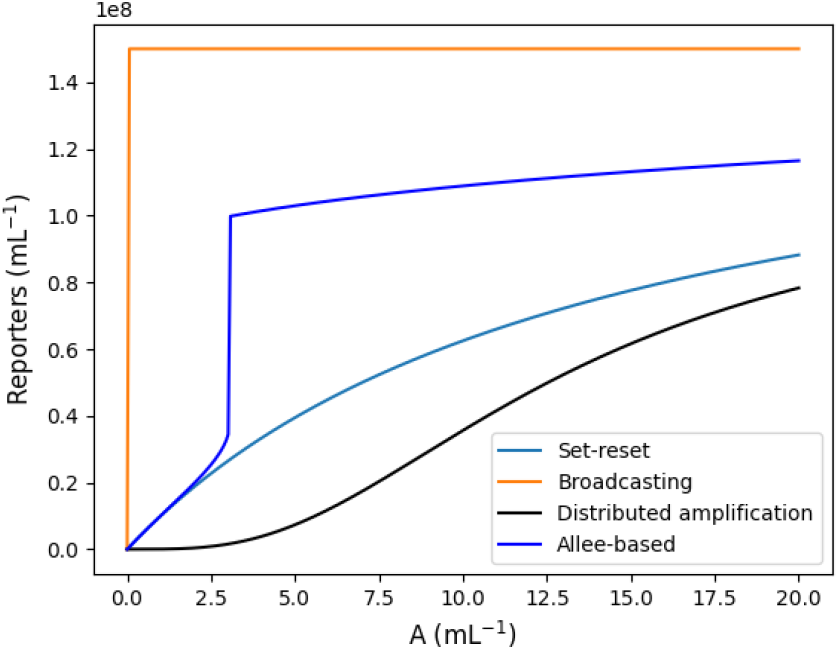
Comparison of the four algorithms: Allee-based, Set-reset, broadcasting, and distributed amplification algorithm. The plot shows steady-state reporter densities (*S* respectively *H*) over analyte concentrations *A* obtained via simulation. Simulation parameters: *P* = 1.5 · 10^8^ mL^−1^, *ρ* = 14 h, *κ* = 35 h, *K* = 8 · 10^7^ mL^−1^, and *n* = 4.

## 6 Robustness Analysis

The non-zero thresholding behavior around a critical threshold *σ*_*c*_ as shown in Theorem 1 and demonstrated by simulations in Figure 5 suggests that the Allee-based algorithm is robust to incorrectly detected analytes by L cells (*σ*_err_ *>* 0 in our model). To demonstrate that this is the case, also for stochastically varying incorrect detections as exprected in a real genetic circuit implementation, we ran simulations where we stochastically varied *σ* over time, simulating the effect of a stochastic *σ*_err_ in absence of an anlyte. For any such simulation, an ideal algorithm is expected to keep H cell densities low, and thus not wrongly signal the detection of an analyte.

Simulations were carried out in MobsPy [9] with a simulated time of 1000 h. For the stochastic model of wrongly detected analytes we chose a stochastic birth-death process of H cells with a birth rate *β* of 0.5 h^−1^ and a death rate *γ* of 2 h^−1^. The parameters have been set to the leaky expression rate and the decay rate of LacI [11] to reflect a realistic parameter range for leaky expression resulting in incorrect L to H transitions of a cell. The so-obtained stochastic rates were then fed into a deterministic transient-time simulation of the Allee-based algorithm. Figure 6 shows the resulting densities *H*(*t*) as well as the rate *σ*(*t*) over time *t* for a simulated time of 100 h. We can observe that while the stochastic event detection rate is capable of increasing the density *H*(*t*), the algorithm does not amplify the H cells further, thus preventing the cells from incorrectly detecting the analyte.

**Figure 6:**
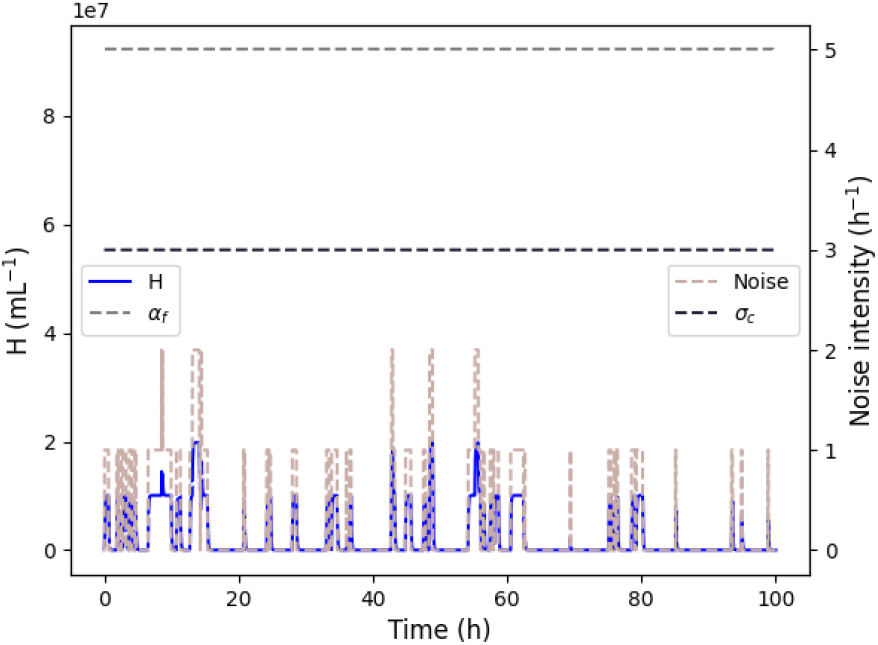
Robustness of the Allee-based algorithm to an incorrect detection of the analyte. The plot shows *H*(*t*) and *σ*(*t*) over time *t* for 4 h simulated time with a low, stochastic rate *σ* in absence of an analyte. The algorithm is seen to not incorrectly amplify the detection of the analyte: the H cell density *H* remains low throughout the simulation. Simulation parameters: *P* = 1.5 · 10^8^ mL^−1^, *ρ* = 14 h, *κ* = 35 h, *K* = 8 · 10^7^ mL^−1^, *n* = 4, *β* = 0.5, and *γ* = 2. Te parameters *α*_*f*_ (see Theorem 1) and *σ*_*c*_ are shown as horizontal lines.

To examine the impact of parameters like the population density *P*, as well as the rate parameter *κ* from (Hold) and *ρ* from (Reset), on the algorithm’s critical threshold *σ*_*c*_, we determined *σ*_*c*_ for parameter sweeps (Figure 7). The parameter ranges that violate the condition of Theorem 1 are marked with setting *σ*_*c*_ = 0 in the heat map.

**Figure 7:**
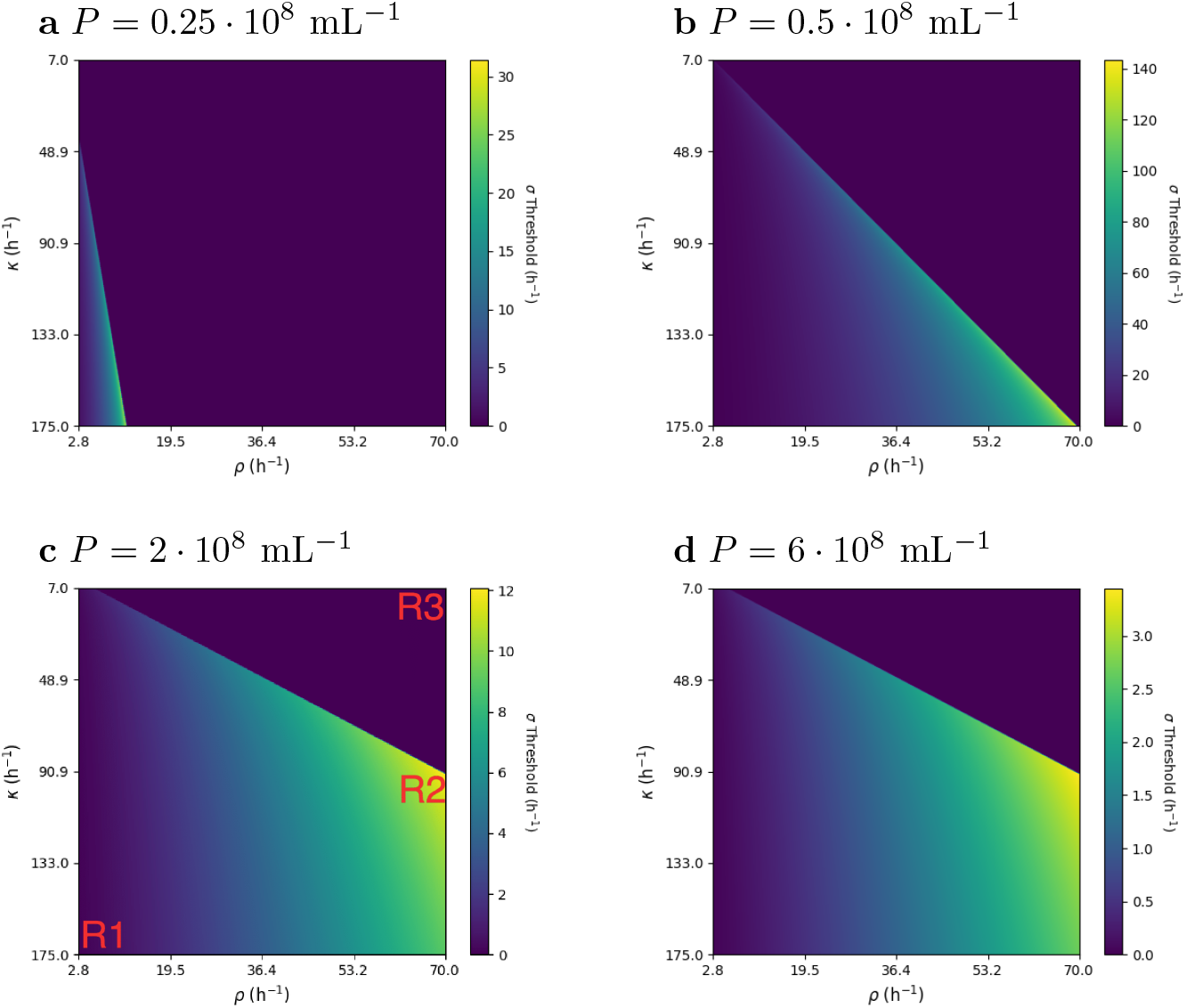
Parameter variations showing *σ*_*c*_ for different reaction rate parameters *κ* and *ρ*. Subplots (a)–(d) differ in the total population density *P*.

In all the heat maps, we can observe a distinct linear boundary separating a region where *σ*_*c*_ = 0 indicating a violation of the condition of 1 and a valid region. For instance, in Figure 7c, the region marked as *R*3 falls in this category where the reset rate constant *ρ* is significantly larger than the hold rate constant *κ*, leading to the inability of the amplification process to trigger. Consequently, the region possesses only a stable equilibrium state with *H* = 0 and no stable equilibrium state for *H* with a high cell density. This is expected as the (Reset) reaction dominates the system behavior in this case.

The heat maps also reveal that when the (Hold) rate parameter *κ* is higher than the (Reset) rate constant *ρ*, the critical threshold *σ*_*c*_ is often close to zero. This is evident in the region marked as *R*1 in Figure 7c. Due to the higher hold rate, the algorithm can produces wrong positives for a lower *σ*.

## 7 Discussion

We presented and discussed a distributed algorithm to detect rare events, such as the presence of a rare analyte, by a population of engineered cells. The algorithm is intended to be used in combination with sensory and reporting circuits within microbial whole-cell sensors. The algorithm is inspired by the Allee effect observed in natural systems: it uses the fact that a certain critical threshold cell density that signal the presence of an analyte is hard to reach initially, but once it is reached, the fact that an analyte has been detected it quickly propagated to the whole cell population.

We have established conditions under which the algorithm provably works as intended (Theorem 1). Numerical simulations of a proof-of-concept circuit demonstrate that the algorithm shows strong amplification of near a critical threshold concentration of the analyte (Figure 5). This is in contrast to three other natural algorithms that have been discussed in this work (Figure 5). Additionally, hybrid stochastic–deterministic simulations (Figure 6) were carried out to demonstrate the robustness of the algorithm to cells that wrongly detect the analyte.

Since the detection thresholds and the targeted total cell populations may differ significantly per application, one may need to adapt the reaction parameters for these cases. We speculate that the simplicity of the detection algorithm as well as the mechanistic understanding of all parameters greatly simplifies this adaption, e.g., via plasmid copy number manipulation to affect the reaction rates [22] and gene removal to alter the parameter *K* [21].

Finally, as has been show by the robustness simulations, the choice of the critical threshold rate *σ*_*c*_ balances the capability of sensing rare events and robustness: while a low threshold favors early detection, a high threshold tolerates a larger concentration of wrongly detected analytes.

## References

[1] WC Allee and Edith S Bowen. Studies in animal aggregations: mass protection against colloidal silver among goldfishes. Journal of Experimental Zoology, 61(2):185–207, 1932.

[2] Silvana Andreescu and Omowunmi A Sadik. Trends and challenges in biochemical sensors for clinical and environmental monitoring. Pure and applied chemistry, 76(4):861–878, 2004.

[3] Philippe Anker, Hugh Mulcahy, Xu Qi Chen, and Maurice Stroun. Detection of circulating tumour dna in the blood (plasma/serum) of cancer patients. Cancer and Metastasis Reviews, 18(1):65–73, 1999.

[4] Shimshon Belkin. Microbial whole-cell sensing systems of environmental pollutants. Current opinion in microbiology, 6(3):206–212, 2003.

[5] Luděk Berec, Elena Angulo, and Franck Courchamp. Multiple allee effects and population management. Trends in Ecology & Evolution, 22(4):185–191, 2007.

[6] Sheng Cai, Yifei Shen, Yan Zou, Peiqing Sun, Wei Wei, Jing Zhao, and Chuan Zhang. Engineering highly sensitive whole-cell mercury biosensors based on positive feedback loops from quorum-sensing systems. Analyst, 143(3):630–634, 2018.

[7] TH Clutton-Brock, D Gaynor, GM McIlrath, ADC Maccoll, R Kansky, P Chadwick, M Manser, JD Skinner, and PNM Brotherton. Predation, group size and mortality in a cooperative mongoose, suricata suricatta. Journal of Animal Ecology, 68(4):672–683, 1999.

[8] Franck Courchamp, Ludek Berec, and Joanna Gascoigne. Allee effects in ecology and conservation. OUP Oxford, 2008.

[9] Fabricio Cravo, Matthias Fuegger, Thomas Nowak, and Gayathri Prakash. Mobspy: A metaspecies language for chemical reaction networks. bioRxiv, 2022.

[10] Tal Danino, Octavio Mondragón-Palomino, Lev Tsimring, and Jeff Hasty. A synchronized quorum of genetic clocks. Nature, 463(7279):326–330, 2010.

[11] M Omar Din, Tal Danino, Arthur Prindle, Matt Skalak, Jangir Selimkhanov, Kaitlin Allen, Ellixis Julio, Eta Atolia, Lev S Tsimring, Sangeeta N Bhatia, et al. Synchronized cycles of bacterial lysis for in vivo delivery. Nature, 536(7614):81–85, 2016.

[12] Yi-Hu Dong and Lian-Hui Zhang. Quorum sensing and quorum-quenching enzymes. The Journal of Microbiology, 43(1):101–109, 2005.

[13] Jianwei He, Xiaoyan Zhang, Yuanyi Qian, Qiyao Wang, and Yunpeng Bai. An engineered quorumsensing-based whole-cell biosensor for active degradation of organophosphates. Biosensors and Bioelectronics, 206:114085, 2022.

[14] Chih-Yuan Hsu, Bing-Kun Chen, Rei-Hsing Hu, and Bor-Sen Chen. Systematic design of a quorum sensing-based biosensor for enhanced detection of metal ion in escherichia coli. IEEE Transactions on Biomedical Circuits and Systems, 10(3):593–601, 2016.

[15] Javier Macía, Francesc Posas, and Ricard V Solé. Distributed computation: the new wave of synthetic biology devices. Trends in biotechnology, 30(6):342–349, 2012.

[16] Melissa B Miller and Bonnie L Bassler. Quorum sensing in bacteria. Annual Reviews in Microbiology, 55(1):165–199, 2001.

[17] Michael S Mooring, Thomas A Fitzpatrick, Tara T Nishihira, and Dominic D Reisig. Vigilance, predation risk, and the allee effect in desert bighorn sheep. The Journal of Wildlife Management, 68(3):519–532, 2004.

[18] Michael Moraskie, Md Harun Or Roshid, Gregory O’Connor, Emre Dikici, Jean-Marc Zingg, Sapna Deo, and Sylvia Daunert. Microbial whole-cell biosensors: Current applications, challenges, and future perspectives. Biosensors and Bioelectronics, 191:113359, 2021.

[19] Lu Pu, Shuai Yang, Aiguo Xia, and Fan Jin. Optogenetics manipulation enables prevention of biofilm formation of engineered pseudomonas aeruginosa on surfaces. ACS synthetic biology, 7(1):200–208, 2018.

[20] David Sidransky. Emerging molecular markers of cancer. Nature Reviews Cancer, 2(3):210–219, 2002.

[21] Parker Smith and Martin Schuster. Antiactivators prevent self-sensing in pseudomonas aeruginosa quorum sensing. Proceedings of the National Academy of Sciences, 119(25):e2201242119, 2022.

[22] Kwei-Lan Tsao and David S Waugh. Balancing the production of two recombinant proteins inescherichia coliby manipulating plasmid copy number: High-level expression of heterodimeric ras farnesyltransferase. Protein expression and purification, 11(3):233–240, 1997.

[23] Janni Vestergaard, Mikkel W Pedersen, Nina Pedersen, Christian Ensinger, Zeynep Tümer, Niels Tommerup, Hans Skovgaard Poulsen, and Lars Allan Larsen. Hedgehog signaling in small-cell lung cancer: frequent in vivo but a rare event in vitro. Lung cancer, 52(3):281–290, 2006.

[24] Seung-Gyun Woo, Sung-Je Moon, Seong Keun Kim, Tae Hyun Kim, Hyun Seung Lim, Gun-Hwi Yeon, Bong Hyun Sung, Chul-Ho Lee, Seung-Goo Lee, Jung Hwan Hwang, et al. A designed whole-cell biosensor for live diagnosis of gut inflammation through nitrate sensing. Biosensors and Bioelectronics, 168:112523, 2020.

[25] Ying Wu, Chien-Wei Wang, Dong Wang, and Na Wei. A whole-cell biosensor for point-of-care detection of waterborne bacterial pathogens. ACS synthetic biology, 10(2):333–344, 2021.

[26] Ludovic Zimmerlin, Vera S Donnenberg, and Albert D Donnenberg. Rare event detection and analysis in flow cytometry: bone marrow mesenchymal stem cells, breast cancer stem/progenitor cells in malignant effusions, and pericytes in disaggregated adipose tissue. In Flow cytometry protocols, pages 251–273. Springer, 2011.

