## Supplementary Material for "An Allee-based Distributed Algorithm for Microbial Whole-Cell Sensors"

### A Mathematical Proofs

#### A.1 Proof of Theorem 1

We start this section with a recall of the model parameters of reactions (Set), (Hold), and (Reset). Let the rare event detection rate $\sigma$ be in $\mathbb{R}_+$, the reset rate constant $\rho$ be in $\mathbb{R}_{>0}$, and the hold rate parameter $\kappa$ be in $\mathbb{R}_{>0}$. Let the maximum population density $P$ be in $\mathbb{R}_{>0}$. Denoting by $H \in \mathbb{R}_+$ the density of H cells, its behavior over time $t \in \mathbb{R}_+$ is given by

$$\frac{dH}{dt} = \sigma \cdot (P - H) + \kappa \frac{H^n}{H^n + K^n}(P - H) - \rho H \quad . \tag{1}$$

We refer to the right-hand side of Equation (1) as the function $s_\sigma$ from $\mathbb{R}_+$ to $\mathbb{R}$, emphasizing the fact that it depends on $\sigma$. In particular, $s_\sigma = s_0 + \sigma \cdot (P - H)$.

The equilibrium points of $H(t)$ are the roots of $s_\sigma$. We start the analysis of the roots by establishing conditions for when $s_\sigma$ has a unique local minimum and a unique local maximum. Those will be later on used to establish the requirements on $s_\sigma$ to have two stable equilibrium points. We begin with a technical lemma:

**Lemma 1.** *Let $\Pi(H) = C - aH - bH^n + cH^{n+1}$ with $H \geq 0$, $C > 0$, $a > 0$, $b > 0$, $c > 0$, and $n > 0$. For $H \geq 0$, one of the following is true:*

1. *There exists an interval $I = ]H_{inc}, H_{dec}[$, such that $\Pi(H) < 0$ if and only if $H$ is in $I$. Furthermore, one has $\Pi(H) = 0$ if and only if $H = H_{inc}$ or $H = H_{dec}$.*

2. *The polynomial $\Pi$ is positive with the exception of at most one zero in $[0, +\infty[$.*

*Proof.* It is $\frac{d\Pi}{dH} = -a - nbH^{n-1} + (n+1)cH^n$. Further, if $H > 0$,

$$\frac{d\Pi}{dH} < 0 \Leftrightarrow a + nbH^{n-1} > (n+1)cH^n \Leftrightarrow \frac{a + nbH^{n-1}}{H^n} > (n+1)c \Leftrightarrow \frac{a}{H^n} + \frac{nb}{H} > (n+1)c \tag{2}$$

As $n > 0$, the expression $\frac{a}{H^n} + \frac{nb}{H}$ is strictly decreasing for $H$ within $\mathbb{R}_+$, converges to 0 as $H$ goes to inf, and converges to inf as $H$ goes to 0. Thus, there exists $H_{\mathrm{d3}} > 0$ such that for all $H$ in $]0, H_{\mathrm{d3}}[$, Equation (2) holds, and for all $H$ in $[H_{\mathrm{d3}}, +\infty[$, Equation (2) does not hold. For the remaining value of $H = 0$ that is needed to complete the analysis in the interval $[0, +\infty[$, one has $\frac{d\Pi}{dH}(0) < 0$. It follows that $\frac{d\Pi}{dH}$ is negative in $[0, H_{\mathrm{d3}}[$, positive in $]H_{\mathrm{d3}}, +\infty[$, and zero at $H_{\mathrm{d3}}$. Therefore, $\Pi$ is strictly decreasing for $H$ within $[0, H_{\mathrm{d3}}]$, strictly increasing for $H$ within $[H_{\mathrm{d3}}, +\infty[$, and there is a unique global minimum for $\Pi$ at $H_{\mathrm{d3}}$.

Let $Q$ be the following predicate: There exists $H_{\mathrm{dr}}$ that is in $[0, H_{\mathrm{d3}}]$ such that $\Pi(H_{\mathrm{dr}}) < 0$. To conclude the proof, one can perform a case distinction over $Q$ and it's negation to show that $Q$ implies statement 1 of the lemma and the negation of $Q$ implies statement 2 of the lemma:

**Case $Q$:** Let us assume there exists an $H_{\mathrm{dr}}$ that is in $[0, H_{\mathrm{d3}}]$ such that $\Pi(H_{\mathrm{dr}}) < 0$. As $\Pi$ is continuous for $H$ in $\mathbb{R}_+$, the intermediate value theorem applies. Thus, as $\Pi(0) = K > 0$, $\Pi(H_{\mathrm{dr}}) < 0$, and $\Pi(H_{\mathrm{dr}})$ is strictly decreasing for $H$ within $[0, H_{\mathrm{d3}}]$, one has that $\Pi$ has a unique root $H_{\mathrm{dinc}}$ in $[0, H_{\mathrm{d3}}]$. Also, as $\Pi(H_{\mathrm{dr}}) < 0$, $\lim_{H \to +\infty} \Pi(H) = +\infty$, and $\Pi$ is strictly increasing for $H$ within $[H_{\mathrm{d3}}, +\infty]$, one has that $\Pi$ has another root $H_{\mathrm{ddec}}$ in $]H_{\mathrm{d3}}, +\infty[$. Thus, there exists an interval $I = ]H_{\mathrm{dinc}}, H_{\mathrm{ddec}}[$, such that $\Pi(H) < 0$ for all $H \in I$ and $\Pi(H) > 0$ for $H \in [0, H_{\mathrm{dinc}}[ \cup ]H_{\mathrm{ddec}}, +\infty[$. Let $H_{\mathrm{dinc}} = H_{\mathrm{inc}}$ and $H_{\mathrm{ddec}} = H_{dec}$ for statement 1 to hold.

**Case $\neg Q$:** Otherwise, there does not exist an $H_{\mathrm{dr}}$ in $[0, H_{\mathrm{d3}}]$ such that $\Pi(H_{\mathrm{dr}}) < 0$. Thus, $\Pi(H_{\mathrm{d3}}) \geq 0$. It follows that $\Pi$ is positive in $[0, +\infty[$ with the exception of at most one zero $H_{\mathrm{d3}}$, as $\Pi$ has a unique global minimum at $H_{\mathrm{d3}}$. Thus, statement 2 of the lemma follows.

Since either statement 1 or statement 2 hold, the lemma follows. □

We next provide a definition to distinguish between different types of $s_\sigma$:

**Definition 1.** *We say $s_\sigma$ is well-shaped if it has a unique local minimum at $H = H_{fmin,\sigma}$ in $\mathbb{R}_{>0}$ and a unique local maximum in $\mathbb{R}_{>0}$ at $H = H_{fmax,\sigma}$, with $H_{fmin,\sigma} < H_{fmax,\sigma}$.*

We also provide the first and second derivatives of $s_\sigma$, as they will be used to study the conditions for which $s_\sigma$ is well-shaped. From the definition of $s_\sigma$:

$$s_\sigma(H) = \sigma \cdot (P - H) + \kappa \frac{H^n}{H^n + K^n}(P - H) - \rho H \tag{3}$$

$$\frac{d}{dH} s_\sigma(H) = \kappa \frac{H^{n-1}(nK^nP - nK^nH - K^nH^n - H^{n+1})}{(K^n + H^n)^2} - \sigma - \rho \tag{4}$$

$$\frac{d^2}{dH^2} s_\sigma(H) = \kappa \frac{nK^nH^{n-2}(nPK^n - PK^n - (nK^n + K^n)H - (nP + P)H^n + (n-1)H^{n+1})}{(K^n + H^n)^3} \tag{5}$$

We now prove a lemma that describes the progression of $\frac{ds_\sigma}{dH}$ as $H$ increases.

**Lemma 2.** *Using the notation from Lemma 1, the derivative $\frac{ds_\sigma}{dH}(H)$ is strictly increasing for $H$ within $[0, H_{inc}]$, strictly decreasing for $H$ within $[H_{inc}, H_{dec}]$, and strictly increasing for $H$ within $[H_{dec}, +\infty[$.*

*Proof.* From Equation (4), since $nK^nH^{n-2} > 0$ for $H > 0$, the sign of $\Pi(H) = nPK^n - PK^n - (nK^n + K^n)H - (nP + P)H^n + (n-1)H^{n+1}$ in any interval $I_\mathrm{d}$ determines if $\frac{ds_\sigma}{dH}$ is increasing, decreasing, or neither for $H$ in $I_\mathrm{d}$.

From Lemma 1 applied to polynomial $\Pi$, either the lemma's statement 1 or 2 holds. Assume by means of contradiction that statement 2 holds. Then, $\frac{ds_\sigma}{dH}$ is strictly increasing for $H$ within $[0, +\infty[$. From the definition of $s_\sigma$, one has $\frac{ds_\sigma}{dH}(0) = -\sigma - \rho > -\kappa - \sigma - \rho$. However, $\lim_{H \to +\infty} \frac{ds_\sigma}{dH} = -\kappa - \sigma - \rho$. As $\frac{ds_\sigma}{dH}(0)$ is greater than its limit to infinity, one has that $\frac{ds_\sigma}{dH}$ cannot be strictly increasing in $H$ in $[0, +\infty[$; a contradiction to statement 2 of Lemma 1.

Therefore Lemma 1 statement 1 has to hold; concluding the proof. □

We next establish the conditions for $s_\sigma$ to be well-shaped using the roots of its derivative $\frac{ds_\sigma}{dH}$ in the following lemma:

**Lemma 3.** *The function $s_\sigma$ is either:*

1. *Not well-shaped with $\frac{ds_\sigma}{dH}$ being negative in $\mathbb{R}_+$ except for at most one zero.*

2. *Well-shaped with both an interval $I = ]H_{d1}, H_{d2}[$, such that for all $H$ in $I$, one has $\frac{ds_\sigma}{dH}(H) > 0$, and a union of two intervals $S = [0, H_{d1}[ \cup ]H_{d2}, +\infty[$, such that for all $H$ in $S$, one has $\frac{ds_\sigma}{dH}(H) < 0$. In this case, $H_{fmin,\sigma} = H_{d1}$ and $H_{fmin,\sigma} = H_{d2}$.*

*Proof.* From Lemma 2, one has that $\frac{ds_\sigma}{dH}$ is strictly increasing for $H$ within $[0, H_\mathrm{inc}]$, strictly decreasing for $H$ within $[H_\mathrm{inc}, H_\mathrm{dec}]$, and strictly increasing for $H$ within $[H_\mathrm{dec}, +\infty[$.

By means of contradiction, let us assume that there exists an $H_\mathrm{db}$ in $[H_\mathrm{dec}, +\infty[$ with $\frac{ds_\sigma}{dH}(H_{db}) \geq 0$. As $\frac{ds_\sigma}{dH}$ is strictly increasing for $H$ within $[H_\mathrm{dec}, +\infty[$, if such $H_\mathrm{db}$ exists in this interval, then for any $H > H_\mathrm{db}$, one has $\frac{ds_\sigma}{dH}(H) \geq 0$. However, from the definition of $s_\sigma$, one has $\lim_{H \to +\infty} s_\sigma(H) = -\infty$, thus, the existence of such an $H_\mathrm{db}$ implies a contradiction to the assumption. It follows that for $H$ in $[H_\mathrm{dec}, +\infty[$, one has:

$$\frac{ds_\sigma}{dH}(H) < 0 \tag{6}$$

Let $T$ be the following predicate: There exists $H_{dp}$ in $[0, H_\mathrm{inc}[$ with $\frac{ds_\sigma}{dH}(H_\mathrm{dp}) > 0$. To prove both statements of Lemma 3, one can show that $\neg T$ implies statement 1 to hold while statement 2 not to hold and $T$ implies statement 2 to hold while statement 1 not to hold:

**Case $\neg T$:** Let us assume that for all $H$ in $[0, H_\mathrm{inc}[$, one has $\frac{ds_\sigma}{dH}(H) \leq 0$. Since, from Lemma 2, for all $H$ within $[0, H_\mathrm{inc}[$, the derivative $\frac{ds_\sigma}{dH}$ is strictly increasing, and by assumption $\frac{ds_\sigma}{dH}(H) \leq 0$, one has $\frac{ds_\sigma}{dH}(H) < 0$ for any $H$ in $[0, H_\mathrm{inc}[$. The inequality becomes a strict one because the interval is open: Assume by means of contradiction that there exists an $H_\mathrm{dint}$ in $[0, H_\mathrm{inc}[$ with $\frac{ds_\sigma}{dH}(H_\mathrm{dint}) = 0$, then there also exists an $H_\mathrm{dcont}$ with $H_\mathrm{dint} < H_\mathrm{dcont} < H_\mathrm{inc}$ for which $\frac{ds_\sigma}{dH}(H_{dcont}) > 0$ by the strict monotonicity of $\frac{ds_\sigma}{dH}$; a contradiction to the initial assumption. Moreover, from Lemma 2, one has that $\frac{ds_\sigma}{dH}$ is strictly decreasing for $H$ within $[H_{\text{inc}}, H_{\text{dec}}]$. Thus, one has that $\frac{ds_\sigma}{dH}(H) < 0$ for $H$ in $]H_{\text{inc}}, H_{\text{dec}}]$. From Equation (6), one has that $\frac{ds_\sigma}{dH}(H) < 0$ for $H$ in $]H_{\text{dec}}, +\infty[$. It follows that $\frac{ds_\sigma}{dH}(H) < 0$ for $H$ in $[0, +\infty[$, except for at most one zero at $H_{\text{inc}}$. Consequently, neither a unique local maximum nor a unique local minimum for $s_\sigma$ exist in $\mathbb{R}_{>0}$. By the definition of well-shaped, $s_\sigma$ is not well-shaped.

**Case** $T$**:** Let us assume that there exists an $H_{\text{dp}}$ in $[0, H_{\text{inc}}[$ with $\frac{ds_\sigma}{dH}(H_{dp}) > 0$.

Since $\frac{ds_\sigma}{dH}$ is continuous for $H$ in $\mathbb{R}_+$, one can use the intermediate value theorem. From the definition of $s_\sigma$ and $H_{\text{dp}}$, one has $\frac{ds_\sigma}{dH}(0) < 0$ and $\frac{ds_\sigma}{dH}(H_{\text{dp}}) > 0$. It follows that $\frac{ds_\sigma}{dH}$ has a unique root $H_{\text{d1}}$ in $]0, H_{\text{dp}}[\subset [0, H_{\text{inc}}]$.

Further, from Equation (6), for all $H$ in $[H_{\text{dec}}, +\infty[$, one has $\frac{ds_\sigma}{dH}(H) < 0$. Take any $H_{\text{dfn}}$ from $[H_{\text{dec}}, +\infty[$, it is $\frac{ds_\sigma}{dH}(H_{\text{dfn}}) < 0$. One can apply the intermediate value theorem to conclude that $\frac{ds_\sigma}{dH}$ has at least one root $H_{\text{d2}}$ between $H_{\text{dp}}$ with $\frac{ds_\sigma}{dH}(H_{\text{dp}}) > 0$ and $H_{\text{dfn}}$ with $\frac{ds_\sigma}{dH}(H_{\text{dfn}}) < 0$. From Lemma 2 we have that $\frac{ds_\sigma}{dH}$ is strictly decreasing in the interval $[H_{\text{inc}}, H_{dec}]$, as well as $H_{\text{dp}}$ in $[H_{\text{inc}}, H_{\text{dec}}]$. Also, from Equation (6), for all $H$ in $[H_{\text{dec}}, +\infty[$, it is $\frac{ds_\sigma}{dH}(H) < 0$, and $H_{\text{dfn}}$ in $[H_{\text{dec}}, +\infty[$. It follows that the root $H_{\text{d2}}$ must be unique.

From Equation (6), one has that $\frac{ds_\sigma}{dH}$ cannot have any roots in $[H_{\text{dec}}, +\infty[$. From the continuity of $\frac{ds_\sigma}{dH}$ in $\mathbb{R}_+$ and the uniqueness of the roots, one has $\frac{ds_\sigma}{dH}(H) > 0$ for any $H$ in $]H_{\text{d1}}, H_{\text{d2}}[$, and $\frac{ds_\sigma}{dH}(H) < 0$ for any $H$ in $[0, H_{\text{d1}}[\cup]H_{\text{d2}}, +\infty[$. Since $\frac{ds_\sigma}{dH}$ only has two roots $H_{\text{d1}}$ and $H_{\text{d2}}$, one has that $s_\sigma$ has a unique local minimum $H_{\text{d1}}$ in $\mathbb{R}_{>0}$ and $s_\sigma$ has a unique local maximum $H_{\text{d2}}$ in $\mathbb{R}_{>0}$. One can conclude that $s_\sigma$ is well-shaped with $H_{\text{fmin},\sigma} = H_{\text{d1}}$ and $H_{\text{fmax},\sigma} = H_{\text{d2}}$.

This concludes the proof. □

We are next studying the roots of $s_\sigma$ with the goal to use a Lyapunov stability argument for convergence. First, we establish that all roots are in $[0, P[$.

**Lemma 4.** *The function $s_\sigma$ does not have any roots greater or equal to $P$.*

*Proof.* For all $H \geq P$, one has $-\rho H < 0$, also $\sigma \cdot (P - H) \leq 0$, and $\kappa \frac{H^n}{H^n + K^n}(P - H) \leq 0$. Therefore, $s_\sigma(H) < 0$ and $s_\sigma$ does not have any roots greater or equal to $P$. □

**Lemma 5.** *The local optima of $s_\sigma$, for $H$ in $\mathbb{R}_{>0}$, are within $]0, P[$.*

*Proof.* Since $s_\sigma$ and $\frac{ds_\sigma}{dH}$ are both continuous and differentiable for $H$ in $\mathbb{R}_+$, a necessary condition for a given $H_{\text{lo}}$ to be a local optima in $\mathbb{R}_{>0}$ is $\frac{ds_\sigma}{dH}(H_{\text{lo}}) = 0$. From Equation (4), for any $H \geq P$, one has $\frac{ds_\sigma}{dH}(H) < 0$. Further, from Equation (4), one has $\frac{ds_\sigma}{dH}(0) = -\sigma - \rho$. The lemma's statement follows. □

We will next show in Lemma 6 that $s_\sigma$ has between one and three roots. In case of three roots, we denote them $H_{\text{ieq},\sigma}$, $H_{\text{ceq},\sigma}$, and $H_{\text{feq},\sigma}$, with $H_{\text{ieq},\sigma} < H_{\text{ceq},\sigma} < H_{\text{feq},\sigma}$. If two roots exist, we denote them $H_{\text{ceq},\sigma}$ and $H_{\text{feq},\sigma}$, with $H_{\text{ceq},\sigma} < H_{\text{feq},\sigma}$. In case of a single root, we denote the root by $H_{\text{feq},\sigma}$. We use $\sigma$ as an index to emphasize that the roots' values depend on $\sigma$. Before showing that there are one to three roots, we state a direct consequence of Lemma 4 and Lemma 5:

**Corollary 3.** *All of the following roots $H_{ieq,\sigma}$, $H_{ceq,\sigma}$, $H_{feq,\sigma}$, and local optima $H_{fmim,\sigma}$, $H_{fmax,\sigma}$ are in $[0, P[$ whenever they exist.*

We next establish necessary and sufficient conditions for $s_\sigma$ to have three roots.

**Lemma 6.** *The following holds:*

- *The function $s_\sigma$ has three roots if and only if it is well-shaped, $s_\sigma(H_{fmin,\sigma}) < 0$, and $s_\sigma(H_{fmax,\sigma}) > 0$, with $H_{ieq,\sigma} < H_{fmin,\sigma} < H_{ceq,\sigma} < H_{fmax,\sigma} < H_{feq,\sigma}$.*
- *The function $s_\sigma$ has two roots if and only if it is well-shaped and either $s_\sigma(H_{fmin,\sigma}) = 0$ or $s_\sigma(H_{fmax,\sigma}) = 0$, with $H_{fmin,\sigma} \leq H_{ceq,\sigma} < H_{fmax,\sigma} \leq H_{feq,\sigma}$.*
- *Otherwise, the function $s_\sigma$ has a single root.*

*Proof.* One has that $s_\sigma$ is continuous for $H$ in $\mathbb{R}_+$, so the intermediate value theorem applies. If $s_\sigma$ is well-shaped, Lemma 3 implies that $s_\sigma$ is strictly decreasing for $H$ within $[0, H_{\text{fmin},\sigma}]$, strictly increasing for $H$ within $[H_{fmin,\sigma} H_{\text{fmax},\sigma}]$, and strictly decreasing for $H$ within $[H_{fmax,\sigma}, +\infty[$. It follows that $s_\sigma$ has a

$$\text{unique minimum at } H_{\text{fmin},\sigma} \in [0, H_{\text{fmax},\sigma}] \tag{7}$$

and a

$$\text{unique maximum at } H_{\text{fmax},\sigma} \in [H_{\text{fmin},\sigma}, +\infty[\ . \tag{8}$$

One can distinguish on whether $s_\sigma$ is not well-shaped and, in case it is well-shaped, the values of $s_\sigma$ at $H_{\text{fmin},\sigma}$ and $H_{\text{fmax},\sigma}$:

- *If $s_\sigma$ is not well-shaped*: Lemma 3 statement 1 implies that $s_\sigma$ is strictly decreasing. Since $s_\sigma(0) = \sigma P$ and $s_\sigma(P) = -\rho P$, $s_\sigma$ has at least one root in $[0, P]$ due to the intermediate value theorem. The root in $[0, P]$ must be unique because $s_\sigma$ is strictly decreasing in $[0, \infty[$.
- *If $s_\sigma(H_{fmin,\sigma}) \geq 0$ and $s_\sigma(H_{fmax,\sigma}) < 0$*: this case is a contradiction to the statement that $s_\sigma$ has a unique minimum at $H_{\text{fmin},\sigma}$ in $[0, H_{\text{fmax},\sigma}]$.
- *If $s_\sigma(H_{fmin,\sigma}) \geq 0$ and $s_\sigma(H_{fmax,\sigma}) \geq 0$*: From (7), only $H_{\text{fmin},\sigma}$ can be a root in $[0, H_{\text{fmax},\sigma}]$. Further, $s(H_{\text{fmax},\sigma}) > s(H_{\text{fmin},\sigma}) > 0$. Since $\lim_{H\to+\infty} s_\sigma(H_{\text{fmax},\sigma}) = -\infty$, and $s_\sigma$ is strictly decreasing for $H$ within $[H_{\text{fmax},\sigma}, +\infty[$, the function $s_\sigma$ has only one root in $]H_{\text{fmax},\sigma}, +\infty[$. It follows that $s_\sigma$ has either one root or, if $s_\sigma(H_{\text{fmin},\sigma}) = 0$, two roots.
- *If $s_\sigma(H_{fmin,\sigma}) < 0$ and $s_\sigma(H_{fmax,\sigma}) \geq 0$*: Since $s_\sigma(0) = \sigma P$, it is $s_\sigma(H_{\text{fmin},\sigma}) < 0$, and $s_\sigma$ is strictly decreasing for $H$ within $[0, H_{\text{fmin},\sigma}]$, the function $s_\sigma$ has one root in $[0, H_{\text{fmin},\sigma}[$. From (8), only $H_{\text{fmax},\sigma}$ can be a root in $[H_{\text{fmin},\sigma}, +\infty[$. It follows that $s_\sigma$ has either one root or, if $s_\sigma(H_{\text{fmax},\sigma}) = 0$, two roots.
- *If $s_\sigma(H_{fmin,\sigma}) < 0$ and $s_\sigma(H_{fmax,\sigma}) > 0$*: From the definition of $s_\sigma$, one has $s_\sigma(0) = \sigma P$ and $\lim_{H\to+\infty} s(H) = -\infty$. From the monotocity of the intervals in Lemma 3 statement 2 and the intermediate value theorem, the function $s_\sigma$ has a root in $[0, H_{\text{fmin},\sigma}[$, another in $]H_{\text{fmin},\sigma} H_{\text{fmax},\sigma}[$, and a last one in $]H_{\text{fmax},\sigma}, +\infty[$. □

As a consequence of Lemma 3 and Lemma 6, we get the following corollaries about the sign of $s_\sigma$:

**Corollary 4.** *If $s_\sigma$ has three roots, then*

- *$s_\sigma(H) > 0$ within $[0, H_{ieq,\sigma}[$,*
- *$s_\sigma(H) < 0$ within $]H_{ieq,\sigma}, H_{ceq,\sigma}[$,*
- *$s_\sigma(H) > 0$ within $]H_{ceq,\sigma}, H_{feq,\sigma}[$, and*
- *$s_\sigma(H) < 0$ within $]H_{feq,\sigma}, +\infty[$.*

*Proof.* From Lemma 6, it is $H_{\text{ieq},\sigma} < H_{\text{fmin},\sigma} < H_{\text{ceq},\sigma} < H_{\text{fmax},\sigma} < H_{\text{feq},\sigma}$. The corollary then follows from the requirements for $s_\sigma(H_{\text{fmin},\sigma}) < 0$ and $s_\sigma(H_{\text{fmax},\sigma}) > 0$ from Lemma 6, and $s_\sigma$ being continuous for $H$ in $\mathbb{R}_+$. □

The following lemma establishes how $s_\sigma$ changes for an increasing parameter $\sigma$.

**Lemma 7.** *For $\varepsilon \in \mathbb{R}_{>0}$, one has $s_{\sigma_1+\varepsilon}(H) > s_{\sigma_1}(H)$ for all $H \in [0, P[$.*

*Proof.* Let $\sigma_2 = \sigma_1 + \varepsilon$. For any $H$ in $[0, P[$, let $d(H) = s_{\sigma_2}(H) - s_{\sigma_1}(H)$. It is,

$$\begin{aligned} d(H) &= s_{\sigma_2}(H) - s_{\sigma_1}(H) \\ &= \sigma_2 \cdot (P - H) + \kappa \frac{H^n}{H^n + K^n}(P - H) - \rho H - \sigma_1 \cdot (P - H) - \kappa \frac{H^n}{H^n + K^n}(P - H) + \rho H \\ &= (\sigma_2 - \sigma_1)(P - H)\ . \end{aligned}$$

Since $(P - H) > 0$ for $H \in [0, P[$, one has that $d > 0$ if $\sigma_2 > \sigma_1$ from which the lemma follows. □

The next corollary is a direct consequence of Corollary 3 and Lemma 7.

**Corollary 5.** *Let $H$ be any of $H_{ieq,\sigma}$, $H_{ceq,\sigma}$, $H_{feq,\sigma}$, $H_{fmim,\sigma}$, or $H_{fmax,\sigma}$. For $\varepsilon \in \mathbb{R}_{>0}$, one has $s_{\sigma+\varepsilon}(H) > s_\sigma(H)$.*

In the next two lemmas, we study the impact of $\sigma$ on the roots.

**Lemma 8.** *Let $s_{\sigma_1}$ be a function with three roots. Let $\varepsilon \in \mathbb{R}_{>0}$ and $\sigma_2 = \sigma_1 + \varepsilon$, such that:*

$$\min_{H \in [0, H_{feq,\sigma_1}]} s_{\sigma_1+\varepsilon}(H) < 0$$

*Then, $s_{\sigma_2}$ has three roots as well. Further, for the first two roots, $]H_{ieq,\sigma_2}, H_{ceq,\sigma_2}[ \subset ]H_{ieq,\sigma_1}, H_{ceq,\sigma_1}[$, and for the third root, $H_{feq,\sigma_1} < H_{feq,\sigma_2}$.*

*Proof.* From Lemma 6, as $s_{\sigma_1}$ has three roots, it is well-shaped. Thus, the roots $H_{\mathrm{ieq},\sigma_1}$, $H_{\mathrm{ceq},\sigma_1}$, and $H_{\mathrm{feq},\sigma_1}$ exist. As $[0, H_{\mathrm{feq},\sigma_1}]$ is a compact set and $s_{\sigma_2}$ is continuous in this compact set, there exists a value $H_{\mathrm{dam}}$ that minimizes $s_{\sigma_2}$ in the domain $[0, H_{\mathrm{feq},\sigma_1}]$. Furthermore, one has that

$$\min_{H \in [0, H_{\mathrm{feq},\sigma_1}]} s_{\sigma_2}(H) < 0$$

by assumption. Thus

$$s_{\sigma_2}(H_{\mathrm{dam}}) < 0 \ . \tag{9}$$

From Corollary 5 and the fact that $H_{\mathrm{feq},\sigma_1}$ is a root $s_{\sigma_1}$, one has that $s_{\sigma_2}(H_{\mathrm{feq},\sigma_1}) > s_{\sigma_1}(H_{\mathrm{feq},\sigma_1}) = 0$. From the definition of $s_\sigma$ one has $\lim_{H\to+\infty} s_{\sigma_2} = -\infty$. It follows from $s_{\sigma_2}$ being continuous in $\mathbb{R}_+$ and the intermediate value theorem, that

$$s_{\sigma_2} \text{ has at least one root in } ]H_{feq,\sigma_1}, +\infty[ \ . \tag{10}$$

Since $s_{\sigma_2}(0) = \sigma_2 P > 0$, $s_{\sigma_2}(H_{\mathrm{dam}}) < 0$ from Equation (9), and $s_{\sigma_2}(H_{\mathrm{feq},\sigma_1}) > 0$, with $0 < H_{\mathrm{dam}} < H_{\mathrm{feq},\sigma_1}$,

$$s_{\sigma_2} \text{ must have at least two roots in } [0, H_{\mathrm{feq},\sigma_1}] \ . \tag{11}$$

It follows that $s_{\sigma_2}$ has at least three roots. From Lemma 6, $s_{\sigma_2}$ can have at most three roots. It follows that $s_{\sigma_2}$ has three roots and $s_{\sigma_2}$ is also well-shaped, so $H_{\mathrm{ieq},\sigma_2}$, $H_{\mathrm{ceq},\sigma_2}$, $H_{\mathrm{feq},\sigma_2}$, $H_{\mathrm{fmin},\sigma_2}$, and $H_{\mathrm{fmax},\sigma_2}$ exist.

Corollary 4 and Lemma 7 imply that for all $H$ in $[0, H_{\mathrm{ieq},\sigma_1}] \cup [H_{\mathrm{ceq},\sigma_1}, H_{\mathrm{feq},\sigma_1}]$, one has $s_{\sigma_2}(H) > 0$ as $s_{\sigma_1}(H) \geq 0$. From statements (11) and (10), the function $s_{\sigma_2}$ has two roots in $[0, H_{\mathrm{feq},\sigma_1}]$ as well as one root in $]H_{\mathrm{feq},\sigma_1}, +\infty[$ (in this order). It follows that $]H_{\mathrm{ieq},\sigma_2}, H_{\mathrm{ceq},\sigma_2}[ \subset ]H_{\mathrm{ieq},\sigma_1}, H_{\mathrm{ceq},\sigma_1}[$ and $H_{\mathrm{feq},\sigma_1} < H_{\mathrm{feq},\sigma_2}$. □

We continue with a definition for the parameters related to the necessary conditions for Theorem 1. We say the parameters are *well-adjusted*, if

$$\max_{H\in[0,P]} \left( \kappa \frac{H^n}{H^n + K^n}(P - H) - \rho H \right) > 0 \ . \tag{12}$$

Noting that $s_0$ is obtained by setting $\sigma = 0$ in $s_\sigma$, one then has

$$s_0 = \kappa \frac{H^n}{H^n + K^n}(P - H) - \rho H \ . \tag{13}$$

We now prove a lemma which will be helpful to show that $s_0$ has three roots if the parameters are well-adjusted:

**Lemma 9.** *There exists a non-empty $I = ]0, H_{di}[$, such that for every $H \in I$, one has $s_0(H) < 0$.*

*Proof.* It is $\frac{d}{dH} s_0(H) = \kappa \frac{H^{n-1}(nK^nP - nK^nH - K^nH^n - H^{n+1})}{(K^n+H^n)^2} - \rho$. As $n > 1$, the polynomial $\Pi = H^{n-1}$ has a non-empty interval $I_{\mathrm{d}} = ]0, H_{\mathrm{dhn}}[$, such that if $H$ in $I_{\mathrm{d}}$, one has $\Pi(H) < \frac{\rho}{\kappa n K^n P}$. One can then bound $\frac{d}{dH}s_0(H)$ for $H$ in $I_{\mathrm{d}}$:

$$\begin{aligned}\frac{d}{dH}s_0(H) &< \kappa\frac{\frac{\rho}{\kappa n K^n P}(nK^nP - nK^nH - K^nH^n - H^{n+1})}{(K^n+H^n)^2}\\ &< \kappa\frac{\frac{\rho}{\kappa n K^n P}(nK^nP)}{(K^n+H^n)^2} - \rho\\ &< \left(\frac{1}{(K^n+H^n)^2} - 1\right)\rho\\ &< 0\end{aligned}$$

It follows that for $H \in I_{\mathrm{d}}$, $\frac{d}{dH}s_0(H) < 0$. Since $s_0(H) = 0$, it is $s_0(H) < 0$ for $H$ in $I_{\mathrm{d}}$; showing the lemma with $I = I_{\mathrm{d}}$. □

We next prove that $s_0$ has three roots if the parameters are well-adjusted. This proof allows us to extend the existence of three roots until a critical point using Lemma 8.

**Lemma 10.** *If the parameters are well-adjusted, the function $s_0$ has three roots: The smallest one at $0$, and two non-zero roots at $\alpha_{di}P$ and $\alpha_{df}P$, with $0 < \alpha_{di} < \alpha_{df} < 1$.*

*Proof.* It is:

$$s_0 = \kappa\frac{H^n}{H^n+K^n}(P-H) - \rho H \tag{14}$$

As $s_0(0) = 0$, the function $s_0$ has a root at 0.

From Lemma 9, there is an interval $I = ]0, H_{\mathrm{di}}[$, such that for $H$ in $I$, one has $s_0(H) < 0$. Choosing any $H_{\mathrm{dri}}$ in $I$, one can conclude that

$$s_0(H_{\mathrm{dri}}) < 0 \ . \tag{15}$$

Let $\arg\max_{H\in[0,P]}(\kappa\frac{H^n}{H^n+K^n}(P-H) - \rho H) = H_{\mathrm{dmax}}$. One has that $H_{\mathrm{dmax}}$ is well-defined as $[0,P]$ is a compact set and $s_0$ is continuous in $[0,P]$. From Equation (12), since the parameters are well-adjusted, one has

$$s_0(H_{\mathrm{dmax}}) > 0 \ . \tag{16}$$

Finally, from Lemma 9, since for any $H$ in $I$, one has $s_0(H) < 0$, one can conclude that

$$H_{\mathrm{dmax}} > H_{\mathrm{dri}} \ , \tag{17}$$

as $s(H_{\mathrm{dmax}}) > 0$ so $H_{\mathrm{dmax}}$ must not be in $I$. From Equations (15), (16), (17), and the intermediate value theorem, the function $s_0$ must have at least one root in $]H_{\mathrm{dri}}, H_{\mathrm{dmax}}[$.

From the definition of $s_0$, one has $s_0(P) = -\rho P$, thus

$$s_0(P) < 0 \ . \tag{18}$$

Thus $H_{\mathrm{dmax}} \neq P$ and, together with the definition of $H_{\mathrm{dmax}}$,

$$H_{\mathrm{dmax}} < P \ . \tag{19}$$

Thus, from Equations (16), (18), (19), and the intermediate value theorem, the function $s_0$ must have at least one root in $]H_{\mathrm{dmax}}, P[$.

Since $s_0$ has at least three roots, Lemma 6 guarantees that there can only be three roots. It follows that, for the roots, $H_{\mathrm{ieq},0} = 0$, $H_{\mathrm{ceq},0}$ in $]H_{\mathrm{dri}}, H_{\mathrm{dmax}}[$, and $H_{\mathrm{feq},0}$ in $]H_{\mathrm{dmax}}, P[$. Setting $\alpha_{\mathrm{di}} = H_{\mathrm{ceq},0}/P$ and $\alpha_{\mathrm{df}} = H_{\mathrm{feq},0}/P$ concludes the proof. □

We will now set $\alpha_{\mathrm{i}} = \alpha_{\mathrm{di}}$ and $\alpha_{\mathrm{f}} = \alpha_{\mathrm{df}}$ as defined in Lemma 10. If the parameters are well-adjusted, for any $s_\sigma$, one can determine $\alpha_{\mathrm{i}}$ and $\alpha_{\mathrm{f}}$ by finding the second and third roots of $s_0 = s_\sigma - \sigma\cdot(P-H)$.

We now prove a lemma for the dependency of $H_{\mathrm{fmax},\sigma}$ as $\sigma$ increases:

**Lemma 11.** *Let $\varepsilon \in \mathbb{R}_{>0}$. If $s_{\sigma_1}$ and $s_{\sigma_1+\varepsilon}$ are well-shaped, one has $H_{fmax,\sigma_1+\varepsilon} < H_{fmax,\sigma_1}$.*

*Proof.* Let $\sigma_2 = \sigma_1 + \varepsilon$. Let $h = \kappa \frac{H^n}{H^n+K^n}(P-H)$. From Lemma 3 statement 2, for any $H$ in the set $[0,+\infty[ \setminus ]H_{\text{fmin},\sigma_1}, H_{\text{fmax},\sigma_1}[$, one has $\frac{d}{dH}s_{\sigma_1}(H) \le 0$. Thus, since $\frac{dh}{dH} = \sigma_1 + \rho + \frac{ds_{\sigma_1}}{dH}$ (see Equation (3)), for any $H_{\text{in}}$ in $]H_{\text{fmin},\sigma_1}, H_{\text{fmax},\sigma_1}[$, and for any $H_{\text{out}}$ in $[0, H_{\text{fmin},\sigma_1}] \cup [H_{\text{fmax},\sigma_1}, +\infty[$, one has $\frac{d}{dH}h(H_{\text{in}}) > \frac{d}{dH}h(H_{\text{out}})$. Because $H_{\text{fmax},\sigma_1}$ is a local optimum of $s_{\sigma_1}$ and $\frac{ds_{\sigma_1}}{dH}$ is continuous for $H$ in $\mathbb{R}_+$, one has $\frac{d}{dH}h(H_{\text{fmax},\sigma_1}) = \sigma_1 + \rho$. As $\sigma_2 > \sigma_1$, one has $\sigma_2 + \rho > \sigma_1 + \rho$. Since $s_{\sigma_2}$ is well shaped, $H_{\text{fmax},\sigma_2}$ exists. Because $H_{\text{fmax},\sigma_2}$ is a local optimum of $s_{\sigma_2}$ and $\frac{ds_{\sigma_2}}{dH}$ is continuous for $H$ in $\mathbb{R}_+$, one has $\frac{d}{dH}h(H_{\text{fmax},\sigma_2}) = \sigma_2 + \rho > \frac{d}{dH}h(H_{\text{fmax},\sigma_1})$. Thus, the only possible value for $H_{\text{fmax},\sigma_2}$ such that $\frac{dh}{dH}(H_{\text{fmax},\sigma_2}) = \sigma_2 + \rho$ must be in $]H_{\text{fmin},\sigma_1}, H_{\text{fmax},\sigma_1}[$. □

Now, we prove that if the parameters are well-adjusted, there exists a critical rate $\sigma_c$ such that $s_{\sigma_c}$ has two roots, for any $\sigma < \sigma_c$, the function $s_\sigma$ has one root, and for any $\sigma > \sigma_c$, the function $s_\sigma$ has three roots. Thus, allowing us to prove the $\sigma$ interval described in Theorem 1.

**Lemma 12.** *If the parameters are well-adjusted, there exists a $\sigma_c$ such that $s_{\sigma_c}$ only has two roots, and one of those roots is $H_{fmin,\sigma_c}$. Furthermore, if $0 \le \sigma < \sigma_c$, $s_\sigma$ has three roots. And, if $\sigma > \sigma_c$, $s_\sigma$ has one root.*

*Proof.* For all $\sigma$ in $\mathbb{R}_+$, $s_\sigma$ is continuous for $H$ in $\mathbb{R}_+$.

As the parameters are well-adjusted, Lemma 10 implies that the function $s_\sigma - \sigma \cdot (P-H) = s_0$ has three roots. Thus, Lemma 6 implies that $s_0$ is well-shaped and both $H_{\text{fmin},0}$ and $H_{\text{fmax},0}$ exist. From Lemma 3, $s_0(H)$ is decreasing for $H$ within $[0, H_{\text{fmin},0}]$ and increasing for $H$ within $[H_{\text{fmin},0}, H_{\text{fmax},0}]$. Thus, one has $\arg\min_{H\in[0,H_{\text{fmax},0}]} s_0(H) = H_{\text{fmin},0}$ and, since $s_0$ has three roots, Lemma 6 implies $s_0(H_{\text{fmin},0}) < 0$. Since $s_\sigma(H)$ is continuous for $\sigma$ in $\mathbb{R}_+$, there exists an interval $\Omega = [0, \sigma_{\text{dci}}[$, for which, for $\sigma$ in $\Omega$, one has $\min_{H\in[0,H_{\text{fmax},0}]} s_\sigma(H) < 0$. Let $\sigma_{\text{dc}}$ be the smallest $\sigma_{\text{dci}}$ such that

$$\min_{H\in[0,H_{\text{fmax},0}]} s_{\sigma_{\text{dc}}}(H) = 0 \quad . \tag{20}$$

One has that $\sigma_{\text{dc}}$ is well-defined because: For any compact set interval $I \subset [0,P[$, let $H_{\text{dmi}} = \arg\min_{H\in I} s_{\sigma_{\text{dh}}}$. One has that $H_{\text{dmi}}$ is well-defined as $I$ is a compact set and $s_\sigma$ is continuous for $H$ in $I$. One can verify that there exists $\sigma_{\text{dh}}$ such that for all $H$ in $I$, one has $s_{\sigma_{\text{dh}}}(H) > 0$, through the following inequality:

$$\sigma_{\text{dh}} > \frac{\kappa \frac{H_{\text{dmi}}^n}{K^n + H_{\text{dmi}}^n}(P - H_{\text{dmi}}) - \rho H_{\text{dmi}}}{(P - H_{\text{dmi}})} \tag{21}$$

Thus, the intermediate value theorem assures that $\sigma_{\text{dc}}$ is finite and well-defined.

Let $H_{\text{msc}} = \arg\min_{H\in[0,H_{\text{fmax},0}]} s_{\sigma_{dc}}(H)$. One has that $H_{\text{msc}}$ is well-defined as $s_\sigma$ is continuous for $H$ in $[0, H_{\text{fmax},0}]$.

As $\sigma_{dc} > 0$ and Lemma 6 implies that $s_0(H_{\text{fmax},0}) > 0$, Corollary 5 assures that:

$$s_{\sigma_{dc}}(H_{\text{fmax},0}) > s_0(H_{\text{fmax},0}) > 0 \tag{22}$$

From the definition of $s_{\sigma_c}$, one has

$$\lim_{H\to\infty} s_{\sigma_{\text{dc}}}(H) = -\infty \quad . \tag{23}$$

From Equations (22) and (23) and the intermediate value theorem,

$$\text{there is at least one root for } s_{\sigma_{\text{dc}}} \text{ in } ]H_{\text{fmax},0}, +\infty[ \quad . \tag{24}$$

It follows from (20) and (24) that $s_{\sigma_{\text{dc}}}$ has at least two roots, so Lemma 6 implies that $s_{\sigma_{dc}}$ is well-shaped.

If $s_{\sigma_{dc}}$ is well-shaped, Lemma 3 statement 2 implies that $s_{\sigma_{dc}}$ is decreasing within $[0, H_{\text{fmin},\sigma_{dc}}]$ and increasing within the interval $[H_{\text{fmin},\sigma_c}, H_{\text{fmax},\sigma_{dc}}]$. Thus, $s_{\sigma_c}$ has a unique minimum $H_{\text{fmim},\sigma_{dc}}$ in $[0, H_{\text{fmax},\sigma_{dc}}]$. One can then use equation (20) to deduce that $s_\sigma$ has only one root in $[0, H_{\text{fmax},\sigma_{dc}}]$ and it must be $H_{\text{fmin},\sigma_{dc}}$.

From Lemma 3, one has that $s_{\sigma_{dc}}$ is strictly decreasing within $[H_{\text{fmax},\sigma_{dc}}, +\infty[$. Therefore, for any $H_1$ and $H_2$, with $H_1 < H_2$ in $[H_{\text{fmax},\sigma_{dc}}, +\infty[$, if $s_{\sigma_{dc}}(H_1) > 0$ and $s_{\sigma_{dc}}(H_2) > 0$, $s_{\sigma_{dc}}$ has no roots in $[H_1, H_2]$. Lemma 11 implies that

$$H_{\text{fmax},\sigma_{dc}} < H_{fmax,0} \quad . \tag{25}$$

From Equation (22), one has $s_{\sigma_{dc}}(H_{\text{fmax},0}) > 0$. Thus, $s_{\sigma_{dc}}$ has no roots in $[H_{\text{fmax},\sigma_{dc}}, H_{\text{fmax},0}]$. It follows that $H_{\text{fmim},\sigma_{dc}}$ is the only root $s_{\sigma_{dc}}$ has in $[0, H_{\text{fmax},0}]$.

From Equation (25) and Lemma 3 implication that $s_{\sigma_{dc}}$ is strictly decreasing in $[H_{\text{fmax},\sigma_0}, +\infty[$, one has that $s_{\sigma_{\text{dc}}}$ is strictly decreasing in $[H_{\text{fmax},\sigma_0}, +\infty[ \subset [H_{fmax,\sigma_{dc}}, +\infty[$. Therefore, using Equation (24), one can conclude that $s_{\sigma_{\text{dc}}}$ has only one root in $[H_{\text{fmax},0}, +\infty[$.

One can conclude that $s_{\sigma_{dc}}$ has only two roots. The root $H_{fmin,\sigma_{dc}}$ in $[0, H_{fmax,0}]$ and the root $H_{\text{feq},\sigma_{dc}}$ in $[H_{fmax,0}, +\infty[$.

From the definition of $\sigma_{dc}$, for any $\sigma < \sigma_{dc}$, one has $\min_{H\in[0,H_{\text{fmax},0}]} s_0(H) < 0$. Therefore, as $s_0$ has three roots, Lemma 8 implies that for any $\sigma < \sigma_{\text{dc}}$, the function $s_\sigma$ has three roots.

Choose any $\sigma > \sigma_{\text{dc}}$. We distinguish between $s_\sigma$ being well-shaped or not:

- If $s_\sigma$ is well-shaped for $\sigma$, Lemma 11 implies that $H_{fmax,\sigma} < H_{fmax,\sigma_{\text{dc}}}$. Lemma 7 implies that for any $H$ in $[0, H_{\text{feq},\sigma_{dc}}]$, one has

$$s_\sigma(H) > s_{\sigma_{dc}}(H) \geq 0 \ . \tag{26}$$

  Lemma 6 implies $H_{\text{feq},\sigma_{dc}} > H_{\text{fmax},\sigma_{dc}} > H_{\text{fmax},\sigma}$. Since Lemma 3 implies that $s_\sigma$ is strictly decreasing for values larger than $H_{\text{fmax},\sigma}$, the function $s_\sigma$ can have at most one root greater than

$$H_{\text{fmax},\sigma} < H_{\text{fmax},0} \ . \tag{27}$$

  From Lemma 6, stating that $s_\sigma$ has between one and three roots, and Equations (26) and (27), the function $s_\sigma$ must have exactly one root in $]H_{\text{feq},\sigma_{\text{dc}}}, +\infty[$.

- If $s_\sigma$ is not well-shaped for $\sigma$, then Lemma 6 guarantees it will only have one root.

The lemma's statement follows from setting $\sigma_{\text{c}} = \sigma_{\text{dc}}$. □

We will use the notation of $\sigma_{\text{c}}$ as defined in Lemma 12 in the following lemmas. We now use the Lyapunov stability theorem to prove convergence of $H(t)$ to a root as $t$ approaches $+\infty$.

**Lemma 13.** *If the parameters are well-adjusted and $\sigma$ in $[0, \sigma_c[$:*

- *If $H(0)$ is in $[0, H_{ceq,\sigma}[$, then $H(t)$ converges to $H_{ieq,\sigma}$.*
- *If $H(0)$ is in $]H_{ceq,\sigma}, +\infty[$, then $H(t)$ converges to $H_{feq,\sigma}$.*

*Proof.* Let $H(0)$ in $[0, H_{\text{ceq},\sigma}[$. If the parameters are well adjusted and $\sigma$ in $[0, \sigma_c[$, Lemma 12 assures that $s_\sigma$ has three roots. Thus, one has that the three roots $H_{\text{ieq},\sigma} < H_{\text{ceq},\sigma} < H_{\text{feq},\sigma}$ exist. We choose the Lyapunov function $V_1$ from $\mathbb{R}$ to $\mathbb{R}$ as:

$$V_1(H) = (H_{\text{ieq},\sigma} - H)^2$$
$$\frac{d}{dH}V_1(H) = -2(H_{\text{ieq},\sigma} - H)$$

Let $\Omega = [0, H_{\text{ceq},\sigma}[$. If both statements hold: *(1)* $s_\sigma(H)\frac{d}{dH}V_1(H) < 0$ for all $H$ in $\Omega$ except for a root of $s_\sigma$ denoted by $H_r$ that is in $\Omega$ and *(2)* $H(0)$ is in $\Omega$. The Lyapunov stability condition guarantees $H(t)$ will converge to the root $H_r$ inside $\Omega$.

From Corollary 4, one has that $s_\sigma$ is positive within $[0, H_{\text{ieq},\sigma}[$. From the definition of $V_1$, it is $\frac{dV_1}{dH}(H) < 0$ for $H$ in $[0, H_{\text{ieq},\sigma}[$. Thus, one has $s_\sigma(H)\frac{d}{dH}V_1(H) < 0$ for $H$ in $[0, H_{\text{ieq},\sigma}[$. From Corollary 4, one has that $s_\sigma$ is negative within $]H_{\text{ieq},\sigma}, H_{\text{ceq},\sigma}[$. From the definition of $V$, $\frac{d}{dH}V_1(H) > 0$ for $H$ in $]H_{\text{ieq},\sigma}, H_{\text{ceq},\sigma}[$. Thus, one has $s_\sigma(H)\frac{d}{dH}V_1(H) < 0$ for $H$ in $]H_{\text{ieq},\sigma}, H_{\text{ceq},\sigma}[$. It follows that $s_\sigma(H)\frac{d}{dH}V_1(H) < 0$ for $H$ in $[0, H_{\text{ceq},\sigma}[$ excluding $H_{\text{ieq},\sigma}$, so the Lyapunov stability condition assures $H(t)$ converges to $H_{\text{ieq},\sigma}$.

One can repeat the same argument for the root $H_{\text{feq},\sigma}$ using the Lyapunov function $V_2(H) = (H_{\text{feq},\sigma} - H)^2$, to conclude $H(t)$ converges to $H_{\text{feq},\sigma}$ if $H(0)$ in $]H_{\text{ceq},\sigma}, +\infty[$. □

**Lemma 14.** *If the parameters are well-adjusted and $\sigma$ in $]\sigma_c, +\infty[$: If $H(0)$ in $[0, +\infty[$, then $H(t)$ converges to $H_{feq,\sigma}$*

*Proof.* From Lemma 12, the only root of $s_\sigma$ is $H_{\text{feq},\sigma}$. The function $s_\sigma$ is continuous for $H$ in $\mathbb{R}_+$. One has that $s_\sigma(0) > 0$, and also that $\lim_{H\to+\infty} s_\sigma = -\infty$. From the previous statements, it follows that $s_\sigma(H) > 0$ for $H$ in $[0, H_{\text{feq},\sigma}[$ and $s_\sigma(H) < 0$ for $H$ in $]H_{\text{feq},\sigma}, +\infty[$.

By using the following Lyapunov function $V$ for the interval $[0, +\infty[$

$$\begin{aligned} V(H) &= (H_{\text{feq},\sigma} - H)^2 \\ \frac{d}{dH} V(H) &= -2(H_{\text{feq},\sigma} - H) \end{aligned} \tag{28}$$

one has $s_\sigma(H)\frac{d}{dH}V(H) < 0$ for $H$ in $[0, +\infty[$ except for $H = H_{feq,\sigma}$. Thus, $H(t)$ converges to $H_{\text{feq},\sigma}$. □

The following lemma is similar to Lemma 8, but it removes the constraint requiring $s_\sigma$ to have three roots and it applies only to $H_{\text{feq},\sigma}$.

**Lemma 15.** *Let $\varepsilon > \mathbb{R}_+$. One has $H_{feq,\sigma+\epsilon} > H_{feq,\sigma}$.*

*Proof.* As $s_{\sigma+\varepsilon}$ is continuous for $H$ in $\mathbb{R}_+$, the intermediate value theorem applies. From Corollary 5, one has that $s_{\sigma+\varepsilon}(H_{\text{feq},\sigma}) > s_\sigma(H_{\text{feq},\sigma}) = 0$ (29). From the definition of $s_{\sigma+\varepsilon}$, one has the following: $\lim_{H\to+\infty} s_{\sigma+\varepsilon} = -\infty$ (30). From Equations (29) and (30), and the intermediary value theorem there must be at least one root in $]H_{\text{feq},\sigma}, +\infty[$. Thus, $H_{\text{feq},\sigma+\epsilon} > H_{\text{feq},\sigma}$. □

Towards the goal of proving the converge to the set $[\alpha_f P, P[$ in Theorem 1, we establish a result on the position of $H_{\text{feq},\sigma}$.

**Lemma 16.** *Let the parameters be well-adjusted. Let $\alpha_i$ and $\alpha_f$ be as defined in Lemma 10. Then $H_{feq,\sigma} \in [\alpha_f P, P[$. If $\sigma \in [0, \sigma_c[$, $H_{ieq,\sigma} \in [0, \alpha_i P[$. Thus, the distance between $H_{ieq,\sigma}$ and $H_{feq,\sigma}$ is at least $(\alpha_f - \alpha_i)P$.*

*Proof.* As the parameters are well-adjusted Lemma 10 implies that $s_\sigma - \sigma(P-H) = s_0$ has three roots, so $\alpha_i = \frac{H_{\text{ceq},0}}{P}$ and $\alpha_f = \frac{H_{\text{feq},0}}{P}$ are well-defined. From Lemma 15, one has $H_{\text{feq},\sigma} > H_{\text{feq},0}$. Thus, Lemma 4 implies $H_{\text{feq},\sigma} \in [\alpha_{\text{f}} P, P[$.

If $\sigma \in [0, \sigma_{\text{c}}[$, from Lemma 8 and Lemma 6, $H_{\text{ieq},\sigma} < H_{\text{ceq},\sigma} < H_{\text{ceq},0}$. Thus, $H_{\text{ieq},\sigma} \in [0, \alpha_{\text{i}} P[$.

It follows that $H_{\text{feq},\sigma} - H_{\text{ieq},\sigma} > H_{\text{feq},0} - H_{\text{ceq},0} > (\alpha_2 - \alpha_1)P$; from which the lemma's statement follows. □

Combining Lemmas 13, 14, 15, 16, as well as setting $H_{\text{c},\sigma} = H_{\text{ceq},\sigma}$, the statement of Theorem 1 follows.

#### A.2 Proof of Theorem 2

We start by defining a lower bound function $b_\sigma$ for $s_\sigma$. We use the index $\sigma$ to emphasize that the lower bound function $b_\sigma$ depends on $\sigma$. If the parameters are well-adjusted, we define $M_\sigma = \min_{H\in[0,H_{\text{ceq},0}]} s_\sigma(H)$. We further define the function $b_\sigma$ from $\mathbb{R}_+$ to $\mathbb{R}$ as:

$$b_\sigma(H) = M_\sigma - \frac{M_\sigma}{H_{\text{feq},\sigma}} H \tag{31}$$

As $s_\sigma$ is continuous, the set $[0, H_{\text{ceq}}, 0]$ is a compact set, and Lemma 6 guarantees that $H_{\text{feq}} > 0$ exists, one has that $M_\sigma$ and $b_\sigma$ are well-defined.

We now prove that $b_\sigma(\text{H})$ is indeed a lower bound for $s_\sigma(\text{H})$ for $H$ in $[0, H_{\text{ceq},0}]$.

**Lemma 17.** *Let the parameters be well-adjusted. If $\sigma$ in $]\sigma_c, +\infty[$, for any $H_1$ and $H_2$ in $[0, H_{ceq,0}]$, one has $s_\sigma(H_1) \geq b_\sigma(H_2) > 0$.*

*Proof.* As the parameters are well-adjusted, Lemma 10 implies that $s_0 = s_\sigma - \sigma \cdot (P - H)$ has three roots. Thus, one has that $H_{\text{ieq},0}$, $H_{\text{ceq},0}$, and $H_{\text{feq},0}$ exist, as well as $[0, H_{\text{ceq},0}] \subset [0, H_{\text{feq},0}]$. From Lemma 15, one has that $[0, H_{\text{feq},0}] \subset [0, H_{\text{feq},\sigma}[$. So,

$$H_{\text{ceq},0} < H_{\text{feq},\sigma} \tag{32}$$

and

$$[0, H_{\mathrm{ceq},0}] \subset [0, H_{\mathrm{feq},\sigma}[\ . \tag{33}$$

Further, $s_\sigma$ is continuous for $H$ in $\mathbb{R}_+$. Since $\sigma$ in $]\sigma_\mathrm{c}, \infty[$, Lemma 12 guarantees that $s_\sigma$ has only one root. From the definition of $s_\sigma$, it is $s_\sigma(0) = \sigma P > 0$, and $\lim_{H\to+\infty} s_\sigma = -\infty$. From the three latter statements, one has

$$s_\sigma(H) > 0 \text{ for } H \text{ in } [0, H_{\mathrm{feq},\sigma}[\ . \tag{34}$$

From statements (33) and (34), it follows that $s_\sigma(H) > 0$ for $H$ in $[0, H_{\mathrm{ceq},0}]$.

One has that $b_\sigma(0) = M_\sigma = \min_{H\in[0,H_{\mathrm{ceq},0}]} s_\sigma(H) \leq s_\sigma(H)$ for all $H$ in $[0, H_{\mathrm{ceq},0}]$. From the definition of $b_\sigma$, this function is strictly decreasing for $H$ in $\mathbb{R}$. It follows that the inequality $s_\sigma(H_1) \geq b_\sigma(H_2)$ holds for all $H_1$ and $H_2$ in $[0, H_{\mathrm{ceq},0}]$.

From Equation (32), $H_{\mathrm{ceq},0} < H_{\mathrm{feq},\sigma}$. From Equation (34), one has the following inequality: $b_\sigma(0) = M_\sigma = \min_{H\in[0,H_{\mathrm{ceq},0}]} s_\sigma(H) \leq 0$. By definition, $b_\sigma$ is an affine function positive for any $H < H_{\mathrm{feq},\sigma}$ and negative for $H > H_{\mathrm{feq},\sigma}$. It follows that $b_\sigma(H)$ is positive for $H$ in $[0, H_{\mathrm{ceq},0}]$. □

The next result is folklore for ODEs. We provide a proof for the sake of completeness.

**Lemma 18.** *Let $\frac{dx}{dt} = f(x)$, where $f$ is a function from $\mathbb{R}$ to $\mathbb{R}$. A solution $x(t)$ of this ODE is monotonic.*

*Proof.* Let us assume by contradiction that $x(t)$ is non-monotonic. This implies that there exist times $t_1$ in $\mathbb{R}^+$, $t_2$ in $\mathbb{R}^+$, and $t_\mathrm{m}$ in $\mathbb{R}^+$ with $t_1 < t_\mathrm{m} < t_2$ such that $x(t_1) = x(t_2)$ and $x(t_\mathrm{m}) \neq x(t_2)$. Consider the integral from time $t_1$ to $t_2$:

$$\int_{t_2}^{t_2} f(x)\frac{dx}{dt}dt \tag{35}$$

By integrating through substitution, one has:

$$\int_{x(t_1)}^{x(t_2)} f(z)dz = 0 \tag{36}$$

Which is equal to zero as $x(t_2) = x(t_1)$. However, since $\frac{dx}{dt} = f(x)$, the first integral also yields:

$$\int_{t_2}^{t_2} (f(x))^2 dt = 0 \tag{37}$$

Since $(f(x))^2 \geq 0$ and $f(x)$ has only asymptotic discontinuities if it is not continuous, the value of the integral is only zero if $f(x)$ is zero almost everywhere. This implies that $x(t)$ is constant for $t$ in $[t_1, t_2]$. It follows that $x(t_\mathrm{m}) = x(t_1) = x(t_2)$, which is a contradiction to the assumption that $x(t)$ is non-monotonic. □

**Lemma 19.** *Let the parameters be well adjusted. Let $H_{ceq,0}$ be the second largest root of $s_0$. Let $\sigma$ be in $]\sigma_c, +\infty[$ and assume that for the initial density $H(0) = 0$.*

*Then, there exists a time $t_h$ such that for any $t > t_h$, one has $H(t) \in ]H_{ceq,0}, +\infty[$. Further, $t_h \leq -\ln\left(1 - \frac{H_{ceq,0}}{H_{feq,\sigma}}\right) \cdot \frac{H_{feq,\sigma}}{M_\sigma}$.*

*Proof.* Consider the two ODEs $\frac{dH^*}{dt} = b_\sigma(H^*) = M_\sigma - \frac{M_\sigma}{H_{\mathrm{feq},\sigma}}H^*$ with $H^*(0) = 0$ and $\frac{dH}{dt} = s_\sigma(H)$ with $H(0) = 0$.

From Lemma 17, one has that $s_\sigma = \frac{dH}{dt} > 0$ for $H$ in $[0, H_{\mathrm{ceq},0}]$. Thus, Lemma 18 implies the function $H(t)$ is strictly increasing while $H(t)$ is in $[0, H_{\mathrm{ceq},0}]$. It follows that there exists a finite time interval $I_1 = [0, t_{\mathrm{dh}}]$ such that for any $t$ in $I$, one has $H(t) \in [0, H_{\mathrm{ceq},0}]$ and for any $t$ not in $I$, one has $H(t) \in ]H_{\mathrm{ceq},0}, +\infty[$. We will later show that we may set $t_\mathrm{h} = t_{\mathrm{dh}}$.

From Lemma 17, one has that $b_\sigma(H^*) = \frac{dH^*}{dt} > 0$ for $H^*$ in $[0, H_{\mathrm{ceq},0}]$. Thus, Lemma 18 implies the function $H^*(t)$ is strictly monotonic increasing while $H^*(t)$ is in $[0, H_{\mathrm{ceq},0}]$. It follows that there exists a there must exist a finite time interval $I_2 = [0, t_{\mathrm{db}}]$ for which for any $t$ in $I$, one has $H^*(t) \in [0, H_{\mathrm{ceq},0}]$ and for any $t$ not in $I$, one has $H^*(t) \in ]H_{\mathrm{ceq},0}, +\infty[$.

Lemma 17 guarantees that for any $H$ and $H^*$ in $[0, H_{\text{ceq},0}]$, one has $s_\sigma(H) \geq b_\sigma(H^*)$. Thus, for any $t$ in the intersection $I_{\text{int}} = [0, t_{\text{dh}}] \cap [0, t_{\text{db}}]$, one has that $H(t)$ and $H^*(t)$ are both in $[0, H_{\text{ceq},0}]$. As $H(0) = H^*(0) = 0$, it is for any $t$ in $I_{\text{int}}$:

$$\int_0^t s_\sigma(H(\tau))d\tau \geq \int_0^t b_\sigma(H^*(\tau))d\tau \tag{38}$$

As $b_\sigma$ and $s_\sigma$ are continuous for $H$ and $H^*$ in $\mathbb{R}_+$, they are integrable in any compact set, thus both integrals are well-defined. From the definition of $t_{\text{dh}}$ and $t_{\text{db}}$, it is:

$$H_{ceq,\sigma} = \int_0^{t_{\text{dh}}} s_\sigma(H(\tau))d\tau$$

$$H_{ceq,\sigma} = \int_0^{t_{\text{db}}} b_\sigma(H^*(\tau))d\tau$$

Therefore:

$$\int_0^{t_{\text{dh}}} s_\sigma(H(\tau))d\tau = \int_0^{t_{\text{db}}} b_\sigma(H^*(\tau))d\tau \tag{39}$$

By means of contradiction, let us assume that $t_{\text{dh}} > t_{\text{db}}$. Thus, $[0, t_{\text{db}}] \subset [0, t_{\text{dh}}]$ and $I_{int} = [0, t_{\text{dh}}]$. Lemma 17 implies $s_\sigma(H) > 0$ for $H$ in $[0, H_{\text{ceq},0}]$. Thus, for $t$ in $[0, t_{\text{dh}}]$, one has $H(t)$ in $[0, H_{\text{ceq},0}]$ with $s_\sigma(H(t)) > 0$ and $s_\sigma(H(t))$ is strictly positive in the interval $I_{\text{int}}$ from (39). It follows from the assumption $t_{\text{db}} < t_{\text{dh}}$:

$$\int_0^{t_{\text{db}}} s_\sigma(H(\tau))d\tau < \int_0^{t_{\text{db}}} b_\sigma(H^*(\tau))d\tau \tag{40}$$

Equation (40) contradicts Equation (38). Thus one can conclude that $t_{\text{dh}} \leq t_{\text{db}}$; showing that the lemma's first statement holds for $t_{\text{h}} = t_{\text{dh}}$.

Finally, the ODE $\frac{dH^*}{dt} = M_\sigma - \frac{M_\sigma}{H_{\text{feq},\sigma}} H^* = b_\sigma(H^*)$ with $H^*(0) = 0$ has as unique solution:

$$H^*(t) = H_{\text{feq},\sigma}\left(1 - e^{-\frac{M_\sigma \cdot t}{H_{\text{feq},\sigma}}}\right)$$

Solving for $t_{\text{db}}$ yields $t_{\text{db}} = -\frac{H_{\text{feq},\sigma}}{M_\sigma} \cdot \ln\left(1 - \frac{H_{\text{ceq},0}}{H_{\text{feq},\sigma}}\right)$; from which the lemma's second statement follows. □

With Lemma 19, we provide a minimum time necessary for $H(t)$ to leave the interval $[0, H_{\text{ceq},0}]$. We now prove that if $H(t)$ has left this interval, it will converge to $[\alpha_{\text{f}} P, P[$.

**Lemma 20.** *Let the parameters be well-adjusted. If there exists a time $t_{in}$ such that $H(t_{in})$ is in $[H_{ceq,0}, +\infty[$, then $H(t)$ converges to $H_{feq,\sigma}$ in $[\alpha_f P, P[$.*

*Proof.* Recall that $\frac{dH}{dt} = s_\sigma$. Since $s_\sigma$ does not depend on time $t$, we may apply a time shift and solve the ODE $\frac{dH^s}{dt_s} = s_\sigma$ where $t_s = t - t_{\text{in}}$ with $t_s \geq t_{\text{in}}$ and $H^s(0) = H(t_{\text{in}})$. Further, for all $t$ in $[t_{\text{in}}, +\infty[$, one has $H^s(t - t_{\text{in}}) = H(t)$. Thus, $H^s(t - t_{\text{in}})$ and $H(t)$ converge to the same value as $t$ goes to infinity. One can now apply Theorem 1 using $H(t_{\text{in}})$ as the initial value $H^s(0) = H(t_{\text{in}})$ to obtain convergence. We next distinguish between different cases for $\sigma$:

**Case $\sigma$ in $[0, \sigma_c[$:** Lemma 12 implies that $s_\sigma$ has three roots. Lemma 8 implies that $H_{ceq,\sigma} < H_{ceq,0} < H^s(0) = H(t_{\text{in}})$, thus Theorem 1 implies that $H^s(t)$ converges to $H^s_{\text{feq},\sigma}$ in $[\alpha_f P, P[$.

**Case $\sigma$ in $]\sigma_c, +\infty[$:** As $[H_{\text{ceq},0}, +\infty[ \subset [0, +\infty[$, Theorem 1 implies that $H^s(t)$ converges to $H_{\text{feq},\sigma}$ in $[\alpha_f P, P[$.

**Case $\sigma = \sigma_c$:** Lemma 10 implies that $s_{\sigma_c}$ has two roots, so Lemma 6 implies that $s_{\sigma_c}$ is well-shaped. Thus, one can apply Lemma 11 to conclude $H_{\text{fmax},\sigma_c} < H_{\text{fmax},0}$. Corollaries 4 and 5 imply that $s_{\sigma_c}(H) > s_0(H) \geq 0$ for any $H$ in $[H_{\text{ceq},0}, H_{\text{fmax},0}]$. Since Lemma 3 implies that $s_{\sigma_c}$ is strictly decreasing for $H^s > H_{\text{fmax},\sigma_c}$ and $\lim_{H \to +\infty} s_{\sigma_c}(H) = -\infty$, the function $s_{\sigma_c}$ has only one root in $[H_{\text{fmax},0}, +\infty[ \subset [H_{\text{fmax},\sigma_c}, +\infty[$, which must be $H_{\text{feq},\sigma_c}$. Since $H_{\text{feq},\sigma_c}$ is contained in $[H_{\text{fmax},0}, +\infty[$ and $H_{\text{ceq},0} < H_{\text{fmax},0}$, it is

$$H_{\text{ceq},0} < H_{\text{feq},\sigma_c} \tag{41}$$

It follows that $s_{\sigma_c}(H) > 0$ for $H$ in $[H_{\mathrm{ceq},0}, H_{\mathrm{feq},\sigma_c}[$ and $s_{\sigma_c}(H) < 0$ for $H$ in $[H_{\mathrm{feq},\sigma_c}, +\infty[$.

From Equation (41), the Lyapunov function $V(H) = (H_{\mathrm{feq},\sigma_c} - H)^2$ for $H$ in $[H_{\mathrm{ceq},0}, +\infty[$ fulfills $\frac{d}{dH}V(H) < 0$ for $H$ in $[H_{\mathrm{ceq},0}, H_{\mathrm{feq},\sigma_c}[$ and $\frac{d}{dH}V(H) > 0$ for $H$ in $[H_{\mathrm{feq},\sigma_c}, +\infty[$.

It follows from the product $s_{\sigma_c} \cdot \frac{d}{dH}V(H)$ that $H^s(t)$ converges to $H_{\mathrm{feq},\sigma_c}$.

Finally, Lemma 16 guarantees that $H_{\mathrm{feq},\sigma_c}$ is in $[\alpha_{\mathrm{f}}P, P[$; which concludes the proof. □

With Lemma 20, one observes that the Allee-based algorithm needs to be exposed to a rare event rate for a time at least $\frac{H_{\mathrm{feq},\sigma}}{M_\sigma} \cdot \ln\left(1 - \frac{H_{\mathrm{ceq},0}}{H_{\mathrm{feq},\sigma}}\right)$ to guarantee that the hold functionality triggers. Once $H(t) > H_{\mathrm{ceq},0}$, the H cell density $H(t)$ will converge to $[\alpha_{\mathrm{f}}P, P[$, no matter the value of $\sigma$ after this time.
